# Temporal changes in the microglial proteome of male and female mice after a diffuse brain injury using label-free quantitative proteomics

**DOI:** 10.1101/2022.05.01.490239

**Authors:** Yasmine V. Doust, Aidan Bindoff, Olivia G. Holloway, Richard Wilson, Anna E. King, Jenna M. Ziebell

**Author notes:** Correspondence: Jenna M. Ziebell. These authors have contributed equally to this work and share senior authorship (Anna E. King, Jenna M. Ziebell).

## Abstract

Traumatic brain injury (TBI) triggers neuroinflammatory cascades mediated by microglia, which promotes tissue repair in the short-term. These cascades may exacerbate TBI-induced tissue damage and symptoms in the months to years post-injury. However, the progression of the microglial function across time post-injury and whether this differs between biological sexes is not well understood. In this study, we examined the microglial proteome in the days (3- and 7-days) to 1 month (28 days) after a midline fluid percussion injury (mFPI) in male and female mice using label-free quantitative proteomics. We identified a reduction in microglial proteins involved with clearance of neuronal debris via phagocytosis at 3- and 7-days post-injury. At 28 days post-injury pro-inflammatory proteins were decreased and anti-inflammatory proteins were increased in microglia. These results indicate a reduction in microglial clearance of neuronal debris in the days post-injury with a shift to anti-inflammatory function by 1 month. The changes in the microglial proteome that occurred across time post-injury did not differ between biological sexes. However, we did identify an increase in microglial proteins related to pro-inflammation as well as insulin and estrogen signalling in males compared with female mice that occurred with or without a brain injury. Although microglial response was similar between males and females up to 1 month following TBI, biological sex differences in the basal microglial proteome has implications for the efficacy of treatment strategies targeting the microglial response post-injury.

## Introduction

Traumatic brain injury (TBI) is a neurological condition that can result in clinical symptoms that include cognitive deficits, memory impairments and sensorimotor dysfunction (Galea et al. 2018; de Freitas Cardoso et al. 2019). The severity of TBI can range from mild to severe, where 80% of TBI cases that present to the clinic are mild to moderate; sustained mostly from falls, motor vehicle accidents and assault (Dhandapani et al. 2012; Laskowski, Creed, and Raghupathi 2015). At the time of injury, mechanical damage occurs to the brain due to movement of the brain within the skull which triggers secondary injury cascades that continue for days to months following TBI (Prins et al. 2013). Typically, secondary injury cascades are thought to resolve within 1- to 3-months after a TBI of mild to moderate severity (Losoi et al. 2016; Carroll et al. 2004; Kwok et al. 2008). However, increasing literature has demonstrated that symptoms can persist for months to years following TBI, which can impact quality of life and ability to return to everyday activities (Nelson et al. 2019; van der Naalt et al. 1999; Carroll et al. 2020). It is currently unclear why particular individuals are susceptible to poor behavioural and cognitive outcomes post-TBI, but it has been theorised to be due to differences in the underlying pathophysiology.

Inflammation of the brain, known as neuroinflammation, is a regularly reported feature following TBI in the clinic and laboratory which is mediated by the primary immune cells of the brain called microglia and supported by other glia such as astrocytes (Cao et al. 2012; Johnson et al. 2013; Xue et al. 2021). Microglia maintain brain homeostasis by performing house-keeping functions such as refining neural circuitry as well as surveying for pathogens and cellular debris (Li and Barres 2018; Nimmerjahn, Kirchhoff, and Helmchen 2005; Tremblay et al. 2011). When microglia detect neuronal damage, such as that which occurs following a TBI, they produce inflammatory cytokines in order to recruit additional immune cells and clear debris via engulfment, also known as phagocytosis (Kofler and Wiley 2011; Bachiller et al. 2018; Gehrmann, Matsumoto, and Kreutzberg 1995). A shift from microglial homeostasis is referred to as microglial reactivity, which typically encompasses inflammatory and phagocytic functionality that is associated with morphological changes (Cao et al. 2012; Stratoulias et al. 2019). Homeostatic microglia have a small circular somata and thin, highly branched processes, whilst, when microglia become reactive, the cell body begins to swell and processes reduce in complexity (Ziebell, Adelson, and Lifshitz 2015; Cao et al. 2012). Microglial reactivity is typically a transient response lasting for days to weeks in order to return brain homeostasis (Cao et al. 2012; Li and Barres 2018). However, immunohistochemical and magnetic resonance imaging (MRI) studies of human cerebral tissue have reported microglial reactivity months to years after a TBI, which may exacerbate neuronal injury and, hence, worsen functional outcomes (Johnson et al. 2013; Chaban et al. 2020; Ramlackhansingh et al. 2011; Gentleman et al. 2004).

A number of studies have investigated how microglia respond over time following a TBI in preclinical models (Lafrenaye et al. 2015; Madathil et al. 2018; Cao et al. 2012; Ziebell et al. 2012; Morrison et al. 2017; Loane et al. 2014). These studies suggest that microglial pro- and anti-inflammation and phagocytosis is beneficial for tissue repair in the days post-injury (Lafrenaye et al. 2015; Madathil et al. 2018; Cao et al. 2012; Ziebell et al. 2012; Morrison et al. 2017; Loane et al. 2014). Conversely, microglial pro-inflammation and phagocytosis that is evident months to years after a TBI has been correlated with increasing neuronal damage and behavioural impairments (Loane et al. 2014; Boone et al. 2019; Ritzel et al. 2020; Saba et al. 2021). Regardless of these findings, a distinct pattern of microglial functions over time following a TBI is still not able to be identified. This could be because particular populations of individuals are more vulnerable to developing long-term inflammation after a TBI than others. Majority of the clinical and experimental TBI research has been conducted in males as they are more likely to endure a TBI due to their participation in risk-taking behaviours, such as reckless driving (Mushkudiani et al. 2007; Gupte et al. 2019). However, females also sustain TBI through a variety of mechanisms, including sport and assault (Gupte et al. 2019). The increasing literature that includes female participants indicates that biological sex can impact TBI recovery, where females exhibit more severe and prolonged symptoms compared with males (Mikolić et al. 2021; Gupte et al. 2019). Although the mechanisms that underlie the biological sex differences in recovery following TBI are currently unknown, microglia have been demonstrated to exhibit biological sex differences in the developing, adult and aged brain as well as following TBI (Doran et al. 2019; Villapol, Loane, and Burns 2017; Yanguas-Cas·s 2020; Villa et al. 2018; Guneykaya et al. 2018). However, the findings from the current literature are contradictory, thus, the differences in microglial function in the healthy brain as well as across time post-TBI between males and females remains unclear.

The current study is the first, to our knowledge, that investigates the proteome of microglia isolated via fluorescence-activated cell-sorting (FACS) from the mouse brain of both biological sexes in the days (3- and 7-days) to 1 month (28 days) following a moderate TBI to get an indication of the short- and long-term microglial response. TBI was reproduced in the laboratory using the midline fluid percussion injury (mFPI) model that produces pathology similar to that which we see in human cases after a global or diffuse brain injury with lack of bruising or cavitation (Rowe, Griffiths, and Lifshitz 2016). Identification of microglial pathways after a TBI and potential therapeutic targets may assist the development of more effective interventions that improve functional outcomes in individuals living with a TBI. We hypothesised that microglia become reactive in the short-term following TBI that’s persists up to 1 month post-injury but differs between biological sexes.

## Methods

### Breeding and housing of mice

Mouse care, handling and surgical procedures were conducted under the Australian Code for the care and use of animals for scientific purposes, 8^th^ edition 2013, and the University of Tasmania’s animal ethics, #A17680, #A18226. Male and female heterogenous CX3CR1^GFP^ mice (n = 48; JAX stock #005582; (Jung et al. 2000)) for fluorescent-activated cell-sorting (FACS) and proteomics and C57BL/6J mice (n = 24) for immunohistochemistry were bred at the University of Tasmania’s Cambridge Farm Facility and transported to the Medical Science Precinct animal facility at 10-weeks of age. Animals were housed in optimouse cages with *ad libitum* access to food and water and a maximum of five mice per cage in 12 – hour light/dark cycles. This work is reported according to the ARRIVE (Animal Research: Reporting *in vivo* Experiments) guidelines.

### Midline fluid percussion injury (mFPI)

Mice were either subjected to a moderate midline fluid percussion injury (mFPI) or left as an uninjured control (naïve) and did not experience any surgical procedures. At 12 - 16 weeks of age, before surgical procedures started, animals were allocated randomly into groups of either naïve, 3-, 7- or 28-days post-injury (n = 6 per sex/group) and the investigator conducting the surgery was blinded to the injury groups to avoid bias in surgery outcomes. Craniotomy surgery and midline fluid percussion injury induction was conducted according to previously published methods (Rowe, Griffiths, and Lifshitz 2016). Briefly, mice were anaesthetised using 5% isofluorane at 1 L/min with 100% oxygen for approximately 5 minutes. Buprenorphine (Temgesic, 0.1 mg/kg) analgesic was then administered via subcutaneous injection, followed by the fur being shaved from the scalp and an injection of local anaesthetic (bupivacaine, 0.025 mg/kg). Animals were left for 20 minutes in the holding cage warmed on a heat pad to onboard analgesics and mitigate pain wind-up. Mice were re-anaesthetised with isofluorane (5% at 1 L/min) and secured in a stereotaxic frame using the ear and bite bars on a heat pad which remained at 37 °C for the duration of the surgery with 3% anaesthesia at 0.3 L/min maintained via nose cone. Antiseptic betadine (Sanofi, Australia) was used to clean the surgical site, which was removed with 70% alcohol solution, and eyes were kept moist with ophthalmic ointment. Anaesthesia was monitored via steadiness of breath, heart rate and absence of reflex, including the tail pinch and pedal withdrawal reflex. A midline sagittal incision was made from between the eyes to just behind the ears in order to expose the skull. The fascia was removed delicately and a thin circular disc was shaved from weed trimmer line and secured to the skull along the sagittal suture, halfway between bregma and lambda. The circular disc was used as a guide for the 3 mm trephine attached to a micro hand-drill chuck which was used to perform a craniotomy by interchanging the direction of turning the trephine. Bone debris was washed away using cool artificial cerebrospinal fluid (aCSF) and once the edges of the craniotomy site were extremely thin, the brain was exposed by gently removing the bone flap without disturbing the underlying dura. An injury hub was placed on top of the craniotomy site and secured onto the skull with cyanoacrylate gel to create a seal between the hub and the skull. The exposed skull was covered with methyl methacrylate cement (dental acrylic), the hub was filled with aCSF to keep the brain moist, and mice were removed from anaesthesia and the stereotaxic frame then placed in a pre-warmed holding cage until upright and alert.

Once mice had recovered for approximately 1-hour, they were reanaesthetised using 5% isoflurane at 1 L/min and the injury hub was visually inspected for any obstructions or blood then filled with aCSF. The injury hub on the mouse was attached to the fluid percussion device to create a continuity of fluid between the injury hub and the fluid filled cylinder. As the animal regained the pedal reflex, the pendulum was released at an angle of 14° to produce a pressure pulse between 1.0 – 1.4 atm, resulting in an injury of moderate severity. The presence of apnea and seizures were recorded as well as the time taken for the mouse to regain the righting reflex which was also used as a measure of injury severity (Figure 1). Mice that had a duration of righting reflex suppression outside of the 5 – 10 minutes range were excluded from the study (n = 2 female CX3CR1^GFP^ mice with righting times of 1min 42sec and 1min 36sec). Once the animal was upright, they were reanaesthetised and the injury hub was removed to inspect the craniotomy site for signs of herniation, hematoma and integrity of the dura. After the wound was closed with sutures, the mice were placed in a pre-warmed holding cage for at least 1-hour of recovery before being returned to the home cage and placed back in the colony. Naïve animals did not undergo any surgical procedures. Post-surgical evaluations were conducted twice daily for 3-days, then once a week until endpoint, to assess behaviour, appearance, suture site and weight for any signs of pain or distress.

**Figure 1:**
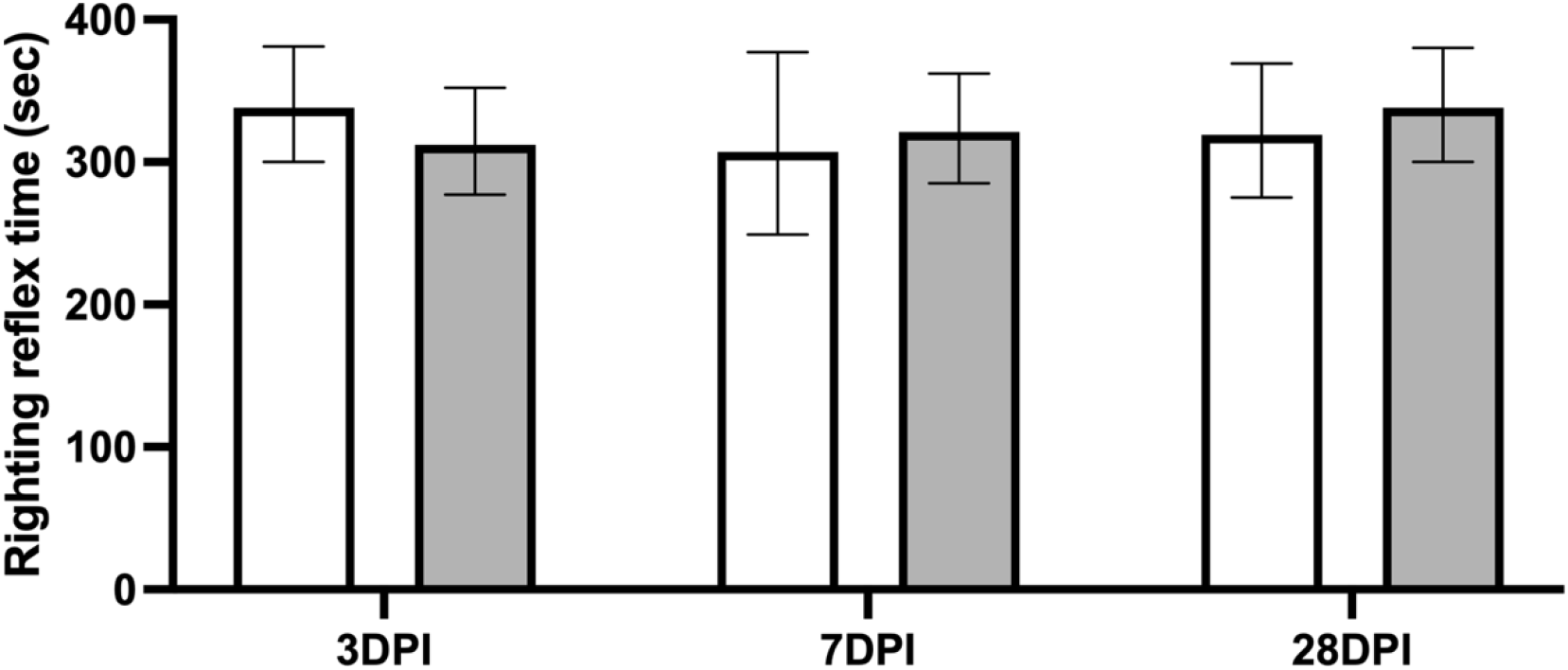
The duration of the righting reflex suppression in female (white) and male (grey) CX3CR1^GFP^ and C57BL/6J mice after a midline fluid percussion injury (mFPI) was used as a measure of injury severity with data represented as estimated marginal means ± 95% confidence intervals (CI). Righting reflex times for each strain were combined for the purpose of this graph. Injury severity was relatively even across the 3-, 7- and 28-days post-injury (DPI) time-points in both biological sexes (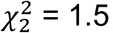; p = 0.4715), where the duration of the righting reflex suppression was between 5 and 10 minutes for all groups which is indicative of a moderate traumatic brain injury (TBI).

### Tissue collection and cell harvesting for proteomics

At 3-, 7- or 28-days post-injury, or in a naïve state, CX3CR1^GFP^ animals were anaesthetised with 5% isoflurane at 1 L/min and given a lethal dose of 115 mg/kg sodium pentobarbitone (Lethobarb, Troy Laboratories, Australia). Mice were transcardially perfused with 0.01M phosphate buffered saline (PBS) and the brain was removed and homogenised through a 70 *μ*m cell strainer (SPL life sciences, cat no. 93070) using a syringe plunger with 20 mL of 0.01M PBS and 0.01 mg/mL of papain (Roche, cat no. 10108014001), 0.05% collagenase (Roche, cat no. 10269638001) and 0.025 U/mL DNAse I (Roche, cat no. 10104159001) into a 50 mL falcon tube (Corning, USA). The 50 mL falcon tube was put into a bead bath at 38°C for 1 hour, inverted every 15 minutes, then centrifuged at 20 RCF at 4°C for 5 minutes. The supernatant was removed and the pellet was triturated with 8 mL of 40% isotonic percoll, containing 1-part 1.5M sodium chloride (NaCl) and 9-parts percoll (Sigma, cat no. E0414-1L), in 0.01M PBS. A 5 mL layer of 0.01M PBS was gently placed on top of the percoll and centrifuged at 50 RCF at 4°C for 45 minutes with no brake. The supernatant was removed that contains myelin and the pellet, containing whole brain cell lysate, was triturated with 2 mL of 0.01M PBS and strained through a 70 *μ*m cell strainer with an extra 3 mL of 0.01M PBS. The sample was then centrifuged at 20 RCF at 4°C for 5 minutes followed by the removal of the supernatant and the pellet triturated in 1 mL of 0.01M PBS in preparation for fluorescence-activated cell sorting (FACS).

### Fluorescence-activated cell sorting (FACS) of microglia

Microglia were isolated from whole brain cell lysate using the BC MoFlo Astrios flow cytometer (Beckman Coulter, USA) which was set up for operation according to the manufacturer’s instructions. Briefly, the stream and lasers were aligned and PMT voltages were confirmed to be optimal using the automated quality control procedure whilst running CST beads. Intellisort was enabled to set up sort parameters, including drop break-off, stream deflection, drop deposition and drop delay. The CyClone robotic arm was configured for sort collection and Summit software was used for obtaining, sorting and interpreting flow cytometry data. The cell lysate was introduced into the Astrios at 60% pressure and microglia were identified as positive for enhanced green fluorescent protein (GFP) under the CX3CR1 promoter. The GFP-positive events, where each event was interpreted as a single microglia, were selected for enrichment sorting via gating and collected in an Eppendorf tube with 100 *μ*L 0.01M PBS. The samples containing approximately 250,000 GFP-positive microglia were centrifuged at 20 RCF at 4°C for 7 minutes. The supernatant was removed followed by the pellet being triturated in 100 *μ*L of lysis buffer (7M Urea, 2M Thiourea, 20mM Tris-HCl) and 1x protease inhibitor (Roche cOmplete mini-inhibitor, cat no. 11836153001) and stored at -80°C prior to protein extraction and digestion.

### Protein extraction and digestion

Microglia were sonicated three times, with 15 seconds in the sonicator and 5 seconds on ice, then kept at 4°C for 2 hours. Protein concentration was determined by performing a Bicinchoninic Acid (BCA) assay as per manufactures protocol (Pierce^TM^ BCA protein assay kit, ThermoScientific) and adjusted to 50 *μ*g/mL. Samples containing ∼5 *μ*g of protein were sequentially reduced then alkylated using standard methods (Hughes et al. 2019), then digested at a 1:25 enzyme:protein ratio with proteomics grade trypsin/rLysC (V5071; Promega, Madison, WI, USA) overnight at 37°C according to the SP3 method (Hughes et al. 2019). Following digestion, peptides were acidified with 2 *μ*l of 1% TFA then desalted using C18 ZipTips® (Millipore, cat no. Z720070-96EA).

### High-performance liquid chromatography and data-independent acquisition (DIA) mass spectrometry (MS)

Peptide samples were randomly allocated into two batches and analysed by DIA-MS using an Ultimate 3000 nano RSLC system coupled with a Q-Exactive HF mass spectrometer fitted with a nanospray Flex ion source (ThermoFisher Scientific, Waltham, MA, USA) and controlled using Xcalibur software (v4.3; ThermoFisher Scientific). Approximately 1 μg of each sample was separated using a 120 minute gradient at a flow rate of 300 nL/min using a 250 mm x 75 μm PepMap 100 C18 analytical column, after preconcentration onto a 20 mm x 75 μm PepMap 100 C18 trapping column. Columns were held at 45°C. MS parameters used for data acquisition were: 2.0 kV spray voltage, S-lens RF level of 60 and heated capillary set to 250°C. MS1 spectra (390 – 1240 m/z) were acquired in profile mode at 120,000 resolution with an AGC target of 3e6. Sequential MS2 scans were acquired across 26 DIA x 25 amu windows over the range of 397.5-1027.5 m/z, with 1 amu overlap between windows. MS2 spectra were acquired in centroid mode at 30,000 resolution using an AGC target of 1e6, maximum IT of 55 ms and normalized collision energy of 27.

### Protein identification, quantification, and normalisation

DIA-MS raw files were processed using Spectronaut software (v14.8; Biognosys AB). A project-specific library was generated using the Pulsar search engine to search the DIA MS2 spectra against the *Mus musculus* UniProt reference proteome (comprising 44,456 entries, April 2017). Biognosys factory settings were used for both spectral library generation and DIA data extraction, with the exception that single-hit proteins were excluded. For library generation, search parameters allowed for N-terminal acetylation and methionine oxidation as variable modifications and cysteine carbamidomethylation as a fixed modification and up to two missed cleavages were permitted. Peptide, protein and PSM thresholds set to 0.01. Mass tolerances were based on first pass calibration and extensive calibration for the calibration and main searches, respectively, with correction factors set to 1 at the MS1 and MS2 levels. Targeted searching of the library based on XIC extraction deployed dynamic retention time alignment with a correction factor of 1. Protein identification deployed a 1% q-value cut-off at precursor and protein levels, automatic generation of mutated peptide decoys based on 10% of the library and dynamic decoy limitation for protein identification. Protein label-free quantitation used the Quant 2.0 setting, based on MS2-level data using the intensity values for the Top3 peptides (stripped sequences) and cross-run normalization was based on median peptide intensity.

### Data filtering and statistical analysis

The Spectronaut protein output table was imported into Perseus software (Tyanova et al., 2016) and proteins identified in fewer than 50% of the samples were excluded from further analysis. Samples that contained less than 1200 proteins were excluded from statistical analysis (n = 2 female mice at 7-days post-injury; n = 2 female mice at 28-days post-injury). Remaining missing values were imputed using default Perseus settings. We used sparse principal components analysis (SPCA) to form a low-dimensional representation of the high-dimensional proteomics matrix. Nominally, these can be interpreted as microglial protein communities. Classical principal components analysis (PCA) is known to be inconsistent when the number of dimensions (in this case proteins, *p*) is greater than the number of observations (in this case samples, *n*). SPCA resolves this issue by imposing a penalty on loadings, such that loadings that contribute little to the component can go to zero, and is consistent when *p >> n*. We used the LASSO penalty (L_1_-norm) on the columns (*p)* and imposed a further constraint that non-zero loadings must be strictly positive. The penalty term was determined by 10-fold cross-validation following the method of (Witten, Tibshirani, and Hastie 2009) using the ‘PM’ package (Witten and Tibshirani 2020) in R (R Core Team 2021). The number of components was selected by trial and error, we stopped at *i* components when none of the explanatory variables were significantly associated with the *i+1_th_* principal component, since these were the variables of interest. The first four principal components explained 38.3% of variance, and there were 1295, 1519, 1234, and 1250 non-zero loadings respectively for each component. The non-zero loadings for each component were then summed to generate a score for each principal component.

Univariate linear regression was used to estimate the association between explanatory variables time post-injury (in days), sex, and the score for each component. All regression models were adjusted to account for batch variation. We standardized each principal component by subtracting each score from the mean (for each component) and dividing by the standard deviation (SD) for each component, so that the effect of each explanatory variable could be interpreted on comparable scales across components. To estimate sex-differences with time post-injury we first fitted models with a Time x Sex interaction term. If the Time x Sex interaction term was not significant (F-statistic using type II sums of squares) we then fitted main effects models (additive terms only) to estimate average differences with time post-injury and sex differences averaged across time post-injury for each component.

Results for the regression analyses are presented as standardized Beta coefficients with 95% confidence intervals (CI) on the SD scale for each component. We considered results significant at *p* < 0.05, and did not adjust for false discovery rate (reducing the number of comparisons by reducing the dimensionality of the data with a penalty prior to analysis, rather than post-hoc).

To aid with visualisation of the SPCA results (Figure 2), we used uniform manifold approximation and projection [UMAP; (McInnes, Healy, and Melville 2018)] to reduce the *p-*dimensional matrix to 2-dimensions. The UMAP algorithm preserves local and global scale and is robust to non-linearities, such that proteins which co-vary across samples are co-located in 2-dimensional space. We shaded points using their corresponding SPCA loadings (unstandardized to preserve zeros), arranged in a panel for each component. We used the ‘umap’ (Konopka 2020) and ‘ggplot2’ (Wickham, Navarro, and Pedersen 2016) packages to produce this figure. All code is available on request.

**Figure 2:**
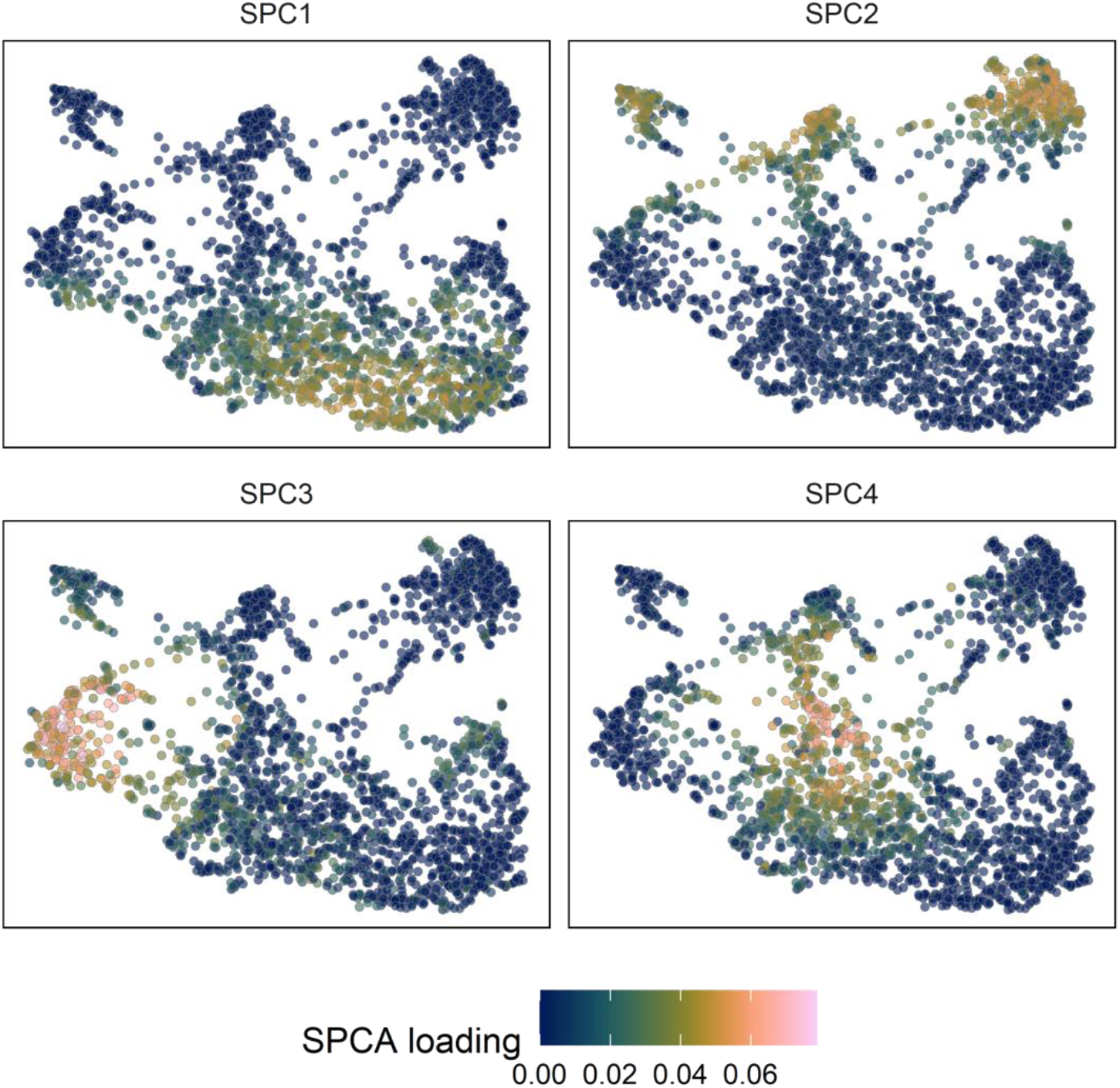
Uniform Manifold Approximation and Projection (UMAP) of all proteins identified within microglia where each dot represents a protein. Protein intensities which were distributed most similarly across samples are co-located on the graph. We have shaded each protein using untransformed loadings from sparse principal components analysis (SPCA), each panel representing one of the first four principal components.

### Bioinformatics

Gene ontology (GO) and WikiPathway analysis was conducted using the STRING database (version 11.5; https://string-db.org/; (Szklarczyk et al. 2018), an online bioinformatic resource. Protein lists included each sparse principal component protein communities which were analysed using the mouse genome database. Over-represented ontologies and pathways were scored with *p*-values <0.05, corrected for multiple testing using the Benjamini-Hochberg procedure, considered statistically significant.

### Tissue collection and preparation for immunohistochemistry

C57BL/6J mice were anaesthetised with isoflurane (5%, 1 L/min) and given a terminal injection of 115 mg/kg sodium pentobarbitone (Lethobarb, Troy Laboratories, Australia) at 3- or 7-days post-injury, or in a naïve state. Once deeply anaesthetised (absence of pedal withdrawal reflex and deep breathing), animals were transcardially perfused using 4% paraformaldehyde (PFA) in 0.01M PBS to clear the vasculature and fix the tissue. The cranium was removed and post-fixed in 4% PFA for 12 hours, then moved to 0.02% sodium azide (NaN_3_) in 0.01M PBS and stored at 4°C prior to sectioning. The brain was removed from the skull in order to perform coronal sectioning at 40 μm using a Leica Microsystems VT1000E vibratome (USA) and the tissue was stored in a 24 well plate in PBS azide at 4°C preceding the conduction of immunohistochemistry.

### Triple-labelling of microglia with phagocytic and myelin markers

Three serial cortical sections (Bregma: 0.74 mm, -1.64 mm, -2.75 mm), per brain, were washed three times in 0.01M PBS for approximately 5 minutes per wash and placed in 100% methanol for 11 minutes at −20°C. Sections were washed again in 0.01M PBS, then placed in 4% normal goat serum in 0.01M PBS blocking solution for 60 minutes with agitation followed by incubation with rat anti-CD68 (1:500; BioRad, cat no. MCA1957GA) and rabbit anti-MBP (1:1000; Sigma, cat no. HPA049222) diluted in 1% block solution and 0.01M PBS overnight at 4°C. All washes and incubations were conducted at room temperature unless otherwise specified. The next day, the tissue sections were washed three times with 0.01M PBS and incubated with goat anti-rat Alexa Fluor 488 (1:1000; Invitrogen, cat no. A-11006) and donkey anti-rabbit Alexa Fluor 594 (1:1000; Invitrogen, cat no. A-21207) diluted in blocking solution for 120 minutes shielded from light. After three 0.01M PBS washes, sections were incubated with rabbit anti-Iba1 red fluorochrome(635)-conjugated antibody (1:250; WAKO, cat no. 013-26471) for 60 minutes at room temperature followed by 7-days at 4°C. Sections were washed with 0.01M PBS and mounted on glass slides (Dako, Denmark) and left to air dry for approximately 10 minutes before being coverslipped (Dako, Denmark) with PermaFluor aqueous mounting medium (Thermo Scientific, cat no. TA-030-FM). Immunohistochemistry experiments were conducted in one batch to avoid batch variation with the inclusion of a negative control (no primary antibody).

### Fluorescent microscopy, manual counts, and statistical analysis of microglia colocalisation

Confocal microscopy was conducted using the Perkin-Elmer UltraVIEW VoX system, involving an inverted Ti Eclipse microscope (Nikon, Japan) with a CSU-X1 spinning disk confocal scanner (Yokogawa Electric Corporation, Japan). The plan apochromatic 40x/0.95 objective (Nikon, Japan) was used via the Volocity 6.3 software to capture z-stack images (1 image taken per 1μm of optical thickness) in the corpus callosum and primary sensory barrel field (S1BF) of both hemispheres with excitation lasers/emission filters of wavelength 488/525, 561/615 and 633/738. Z-stacks were saved as OME.TIFF files for subsequent analysis as maximum intensity projection images that displays the multiple planes of the section in 2 dimensions. Microglial colocalisation with phagocytic (CD68) and myelin (MBP) markers was conducted according to previously published methods (Doust et al. 2021). Briefly, a 1 mm^2^ grid was placed across the image and all microglia were counted within each square, at random, using the cell counter tool, with a target of 100 microglia. The total number of microglia and the number of microglia colocalised with CD68 and MBP were summed for all six images per animal. The proportion of colocalised microglia per animal out of the total number of cells counted was determined by dividing the number of colocalised microglia by the total number of microglia then averaged across each group with 95% confidence intervals (CI). The binomial proportion of total microglia that were colocalised with CD68 and MBP was estimated using logistic regression in a generalised linear mixed effects model (Mcculloch and Neuhaus 2014). Random intercepts were fitted for each animal to account for clustering in the data and *p*-values were computed with likelihood ratio tests using the “lme4” package (Bates et al. 2015) in the R statistical (R Core Team 2021) computing environment. The “emmeans” R package (Lenth et al. 2021) was used to compute estimated marginal means and *post-hoc* contrasts, with corrections for multiple comparisons using Tukey’s method to appropriately control Types 1 errors.

## Results

This study examined the proteins in microglia isolated from male and female CX3CR1^GFP^ mice at 3-, 7- or 28-days after a mFPI as well as those in a naïve state to determine whether the microglial proteome changes with time post-injury or between biological sexes.

### Over 2000 proteins were detected within microglia isolated from the adult mouse brain

Mass spectrometry detected 2,375 proteins and measured the relative intensity or abundance of each protein within approximately 250,000 microglia isolated from male and female mice at 3-, 7- or 28-days post-injury or in a naïve state. Nominal microglial protein communities were identified using sparse principal component analysis (SPCA). Proteins which had a similar pattern of intensity across all samples were grouped into one of the four principal components, where proteins can belong to more than one principal component (Figure 2). Each sparse principal component indicates one protein community that contains proteins that may or may not be related to one another but had a similar pattern of intensity changes across time post-injury or between sexes. SPCA generated a loading value for each protein, where a positive loading value signifies a positive correlation with that principal component. Whilst, a zero value indicates little to no correlation with that principal component. The non-zero loadings were then summed to yield a score for each component.

Principal component scores were compared between samples collected from naïve animals and those at 3-, 7- or 28-days post-injury. Scores from both biological sexes were also compared in a naïve state and at 3-, 7- or 28-days post-injury. Sparse principal component 1 (SPC1; Figure 2) scores were significantly decreased in microglia isolated from mice at 28 days following a mFPI compared with naïve controls (*p* = 0.003; see Table 1; Figure 3). Whilst, SPC2 (Figure 2) scores were significantly increased in microglia isolated from mice at 28 days post-injury compared with naïve animals (*p* = 0.002; see Table 1; Figure 3). Compared to naïve controls, SPC3 (Figure 2) scores were significantly decreased at 3- (*p* < 0.001) and 7-days post-injury (*p* = 0.014) but returned to naïve levels by 28 days post-injury (see Table 1; Figure 3). These differences in SPC1, SPC2 or SPC3 scores with time post-injury were evident regardless of sex (*p* = 0.266, *p* = 0.284, *p* = 0.823, respectively). However, SPC4 (Figure 2) scores were significantly increased in microglia isolated from male mice compared with those isolated from females (*p* = 0.046; see Table 1) which was not dependent upon injury (*p* = 0.322).

**Table 1:** Sparse principal component (SPC) scores with time post-injury, compared with naïve levels, and between sexes (predictors).

| Predictors | SPC 1 |  |  | SPC 2 |  |  | SPC 3 |  |  | SPC 4 |  |  |
| --- | --- | --- | --- | --- | --- | --- | --- | --- | --- | --- | --- | --- |
|  | std. Beta | standardized CI | p | std. Beta | standardized CI | p | std. Beta | standardized CI | p | std. Beta | standardized CI | p |
| (Intercept) | 0.33 | -0.20 – 0.85 | 0.247 | -0.35 | -0.87 – 0.17 | 0.235 | 0.76 | 0.28 – 1.24 | **0.003** | -0.34 | -0.89 – 0.22 | **0.038** |
| Time [3d] | -0.02 | -0.77 – 0.72 | 0.949 | 0.02 | -0.71 – 0.75 | 0.96 | -1.52 | -2.20 – -0.84 | **<0.001** | 0.75 | -0.03 – 1.54 | 0.06 |
| Time [7d] | -0.12 | -0.96 – 0.72 | 0.778 | 0.2 | -0.63 – 1.03 | 0.629 | -0.98 | -1.75 – -0.21 | **0.014** | 0.28 | -0.62 – 1.17 | 0.533 |
| Time [28d] | -1.25 | -2.04 – -0.47 | **0.003** | 1.29 | 0.52 – 2.06 | **0.002** | -0.57 | -1.28 – 0.15 | 0.115 | 0.29 | -0.54 – 1.12 | 0.486 |
| Sex [Male] | 0.08 | -0.21 – 0.37 | 0.595 | -0.1 | -0.39 – 0.18 | 0.474 | -0.25 | -0.51 – 0.02 | 0.064 | 0.31 | 0.01 – 0.62 | **0.046** |
| Batch | -0.14 | -0.42 – 0.15 | 0.344 | 0.14 | -0.14 – 0.42 | 0.328 | 0.01 | -0.25 – 0.27 | 0.925 | 0.16 | -0.14 – 0.46 | 0.288 |
| Observations | 42 |  |  | 42 |  |  | 42 |  |  | 42 |  |  |
| R<sup>2</sup> / R<sup>2</sup><br>adjusted | 0.293 / 0.195 |  |  | 0.313 / 0.218 |  |  | 0.413 / 0.331 |  |  | 0.209 / 0.099 |  |  |

Standardized beta coefficients (unit standard deviation scale) and their 95% CI are shown. Bold p-values indicate that the coefficients are statistically significant.

**Figure 3:**
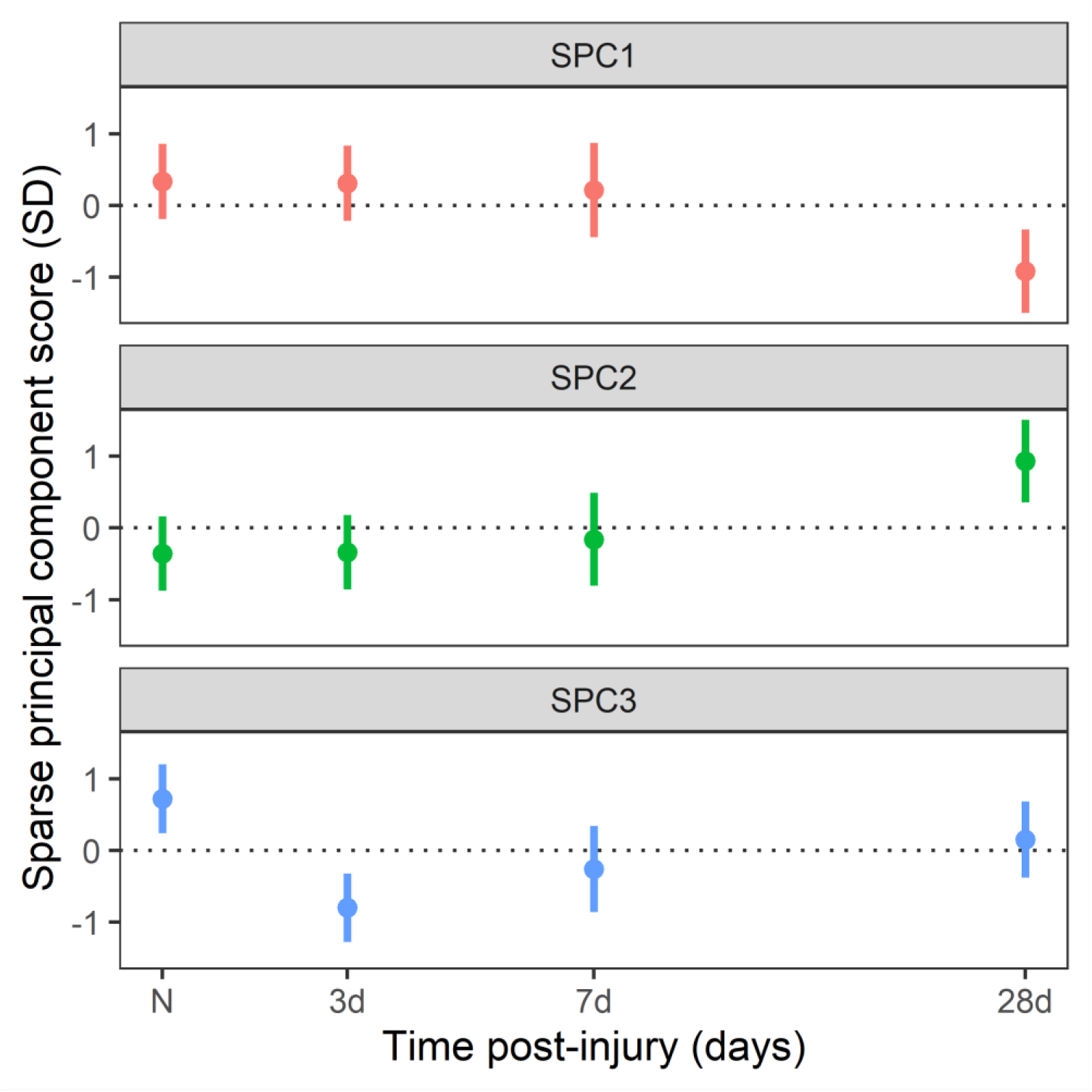
Sparse principal component scores (decomposed using SPCA) across time post-injury (days). The first sparse principal component (SPC) scores significantly decreased from naïve levels at 28 days post-injury, whilst the second SPC scores increased at 28 days post-injury. The third SPC scores significantly decreased at 3- and 7-days post-injury but returned to naïve levels by 28 days post-injury. Estimated means and 95% confidence intervals are shown on unit standard deviation scale.

### Microglia isolated from mice at 3- and 7-days post-injury had a reduction in proteins related to phagocytosis of myelin and synapses that returned to naïve levels by 28-days post-injury

The third SPC scores (Figure 2) were significantly decreased in microglia isolated from mice at 3- and 7-days following mFPI but returned to naïve levels by 28 days post-injury (see Table 1; Figure 3). Using STRING gene ontology (GO) analysis, SPC3 protein communities were identified as being related to synapses and the myelin sheath (Figure 4A). STRING Wikipathway analysis also revealed that SPC3 protein communities were involved with phagocytosis of neuronal debris (“regulation of the actin cytoskeleton:WP523”, “spinal cord injury pathway:WP2432”; Figure 4B). The proteins related to the “regulation of the actin cytoskeleton:WP523” are known to be involved with the formation phagosomes during phagocytosis (Mao and Finnemann 2015). Additionally, proteins related to the “spinal cord injury pathway:WP2432” include myelin proteins (Mbp, Mag; Figure 4B) and FK506 binding protein 1a (Fkbp1a) which is involved in Fc gamma R mediated phagocytosis, suggesting that the proteins related to the “spinal cord injury pathway:WP2432” are involved with myelin phagocytosis (Bohdanowicz and Fairn 2011). SPC3 protein communities were also related to proliferation (“IL-5 signalling pathway:WP151”, “IL-3 signalling pathway:WP373”, “EGFR1 signalling pathway:WP572”) and cell migration (“chemokine signalling pathway:WP2292”; Figure 4B). Proteins related to proliferation and cell migration were also involved with phagocytosis of neuronal debris (see Table 2; Figure 5) indicating that the phagocytosis pathways related to SPC3 protein communities were intermingled with other immune-related functions. SPC3 protein communities were also involved with amino acid metabolism (“amino acid metabolism:WP662”) and glycolysis (“glycolysis and gluconeogenesis:WP157”) as well as oxidative phosphorylation (“oxidative phosphorylation:WP1248”, “electron transport chain:WP295”, “TCA cycle:WP434”; Figure 4B). These results suggest that amino acid metabolism, glycolysis and oxidative phosphorylation are associated with microglial phagocytic activity which is in line with previous literature that have extensively reported that microglia utilise oxidative phosphorylation when undertaking phagocytic functions (reviewed in Lauro and Limatola 2020; Yang et al. 2021). However, glycolysis and amino acid metabolism is typically correlated with pro-inflammatory functions (Geric et al. 2019; reviewed in Yang et al. 2021; Lauro and Limatola 2020). Our findings support the increasing research that suggests that phagocytic microglia adopt a mixed mode of metabolism by utilising glycolysis, amino acid metabolism and oxidative phosphorylation (Wang et al. 2019). As SPC3 scores were significantly decreased from naïve levels in microglia isolated from mice at 3- and 7-days post-injury, these data indicates that microglial phagocytosis of myelin and synapses was reduced at 3- and 7-days post-injury compared with naïve controls. As we hypothesise that myelin was being phagocytosed by microglia, immunohistochemistry (IHC) was used to examine spatial relationships between microglia and engulfment of myelin.

**Figure 4:**
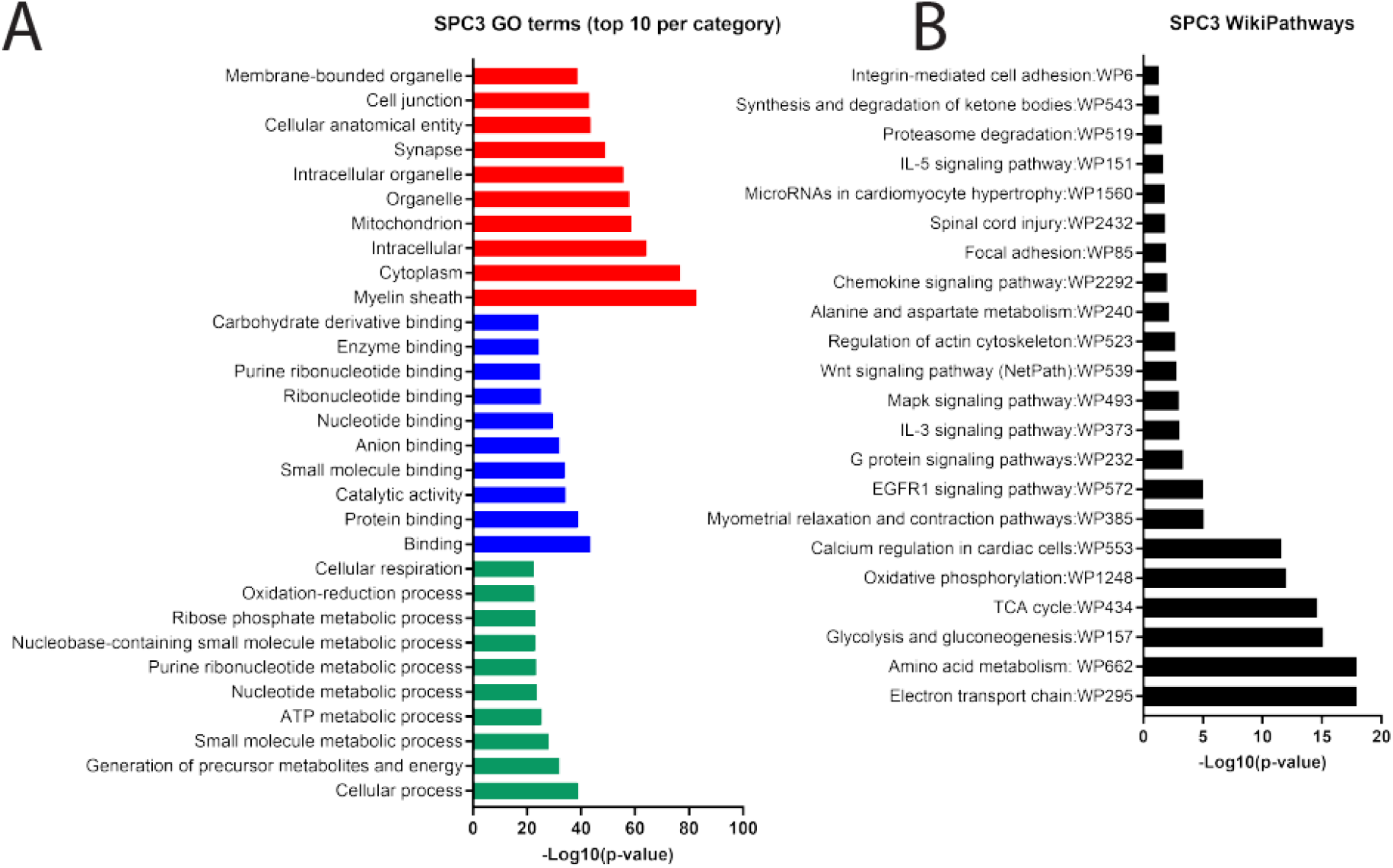
STRING gene ontology (GO; A) and WikiPathway (B) analysis of sparse principal component 3 (SPC3) protein communities. GO analysis includes the top 10 GO terms per category; biological process (green), molecular function (blue) and cellular component (red). Made with GraphPad Prism, version 8.

**Figure 5:**
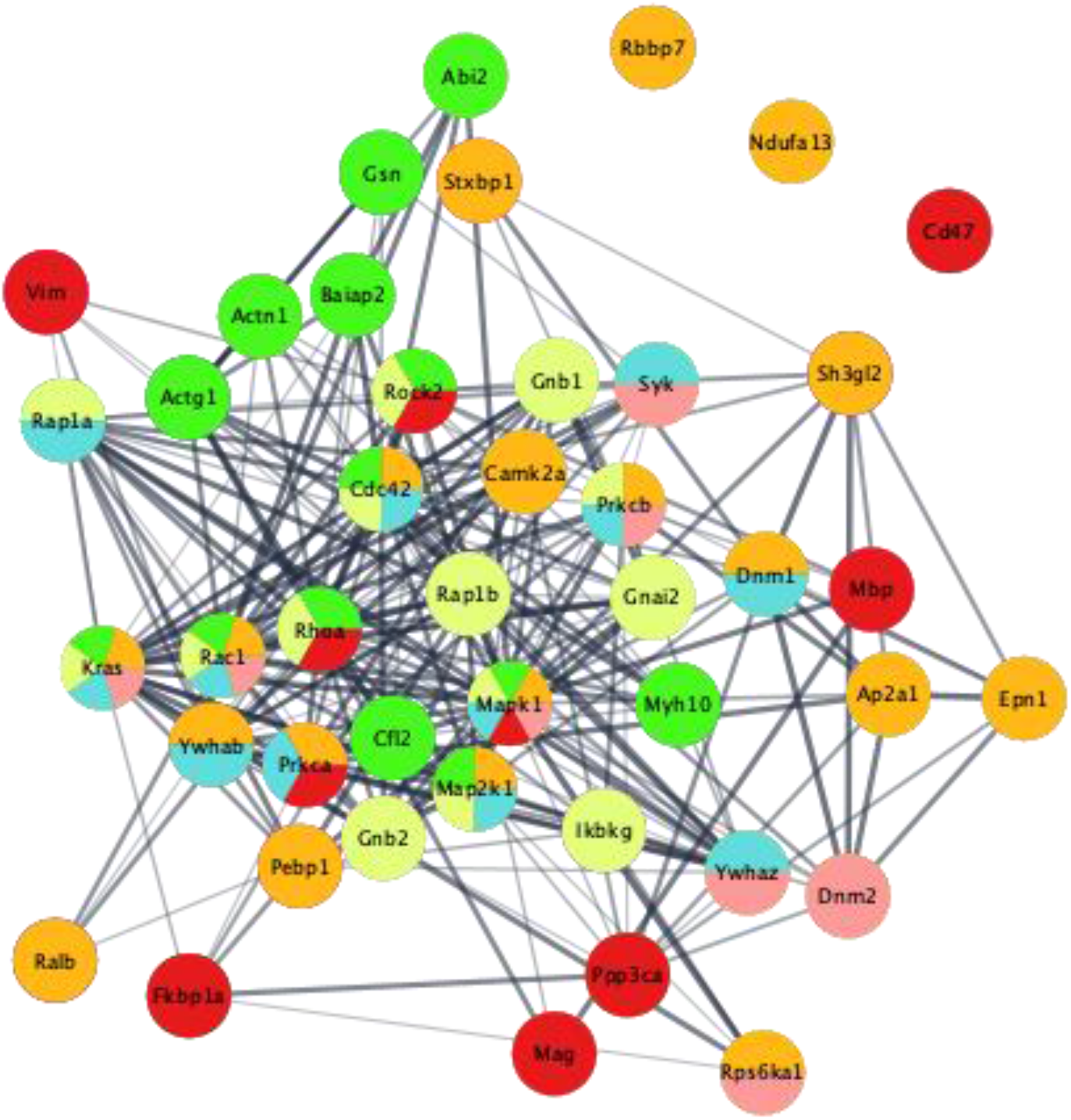
STRING protein network of sparse principal component 3 (SPC3) protein communities involved with proliferation (“EGFR1 signalling:WP572” - orange, “IL-3 signalling:WP373” - blue, “IL-5 signalling” - pink), cell migration (“chemokine signalling:WP2292” - yellow) and myelin phagocytosis (“regulation of actin cytoskeleton:WP523” – green, “spinal cord injury:WP2292” - red) identified via STRING Wikipathway analysis.

**Table 2:** Sparse principal component 3 (SPC3) protein communities that were involved with proliferation (“EGFR1 signalling:WP572”, “IL-3 signalling:WP373”, “IL-5 signalling:WP151”) and cell migration (“chemokine signalling:WP2292”) as well as myelin phagocytosis (“regulation of actin cytoskeleton:WP523”, “spinal cord injury:WP2432”) identified via STRING WikiPathway analysis.

| EGFR1 | IL-3 | Regulation of actin cytoskeleton | Chemokines | IL-5 | Spinal cord injury |
| --- | --- | --- | --- | --- | --- |
| Ralb | Map2k1 | Map2k1 | Map2k1 | Ywhaz | Rhoa |
| Map2k1 | Ywhab | Rhoa | Rhoa | Kras | Rock2 |
| Ywhab | Ywhaz | Rock2 | Rock2 | Mapk1 | Vim |
| Sh3gl2 | Kras | Actn1 | Gnb1 | Prkcb | Fkbp1a |
| Kras | Cdc42 | Baiap2 | Gnb2 | Rac1 | Mbp |
| Rbbp7 | Prkca | Gsn | Kras | Rps6ka1 | Ppp3ca |
| Pebp1 | Mapk1 | Kras | Cdc42 | Syk | Prkca |
| Cdc42 | Prkcb | Cdc42 | Gnai2 | Dnm2 | Mapk1 |
| Prkca | Rac1 | Abi2 | Mapk1 |  | Cd47 |
| Mapk1 | Rap1a | Mapk1 | Rap1b |  | Mag |
| Prkcb | Dnm1 | Actg1 | Prkcb |  |  |
| Rac1 | Syk | Cfl2 | Rac1 |  |  |
| Dnm1 |  | Rac1 | Rap1a |  |  |
| Stxbp1 |  | Myh10 | Ikbkg |  |  |
| Epn1 |  |  |  |  |  |
| Camk2a |  |  |  |  |  |
| Rps6ka1 |  |  |  |  |  |
| Ndufa13 |  |  |  |  |  |
| Ap2a1 |  |  |  |  |  |

Coronal sections from male and female C57BL/6J mice at 3- and 7-days post-injury and naïve animals were stained with ionising calcium binding adaptor protein 1 (Iba1), cluster of differentiation 68 (CD68), and myelin basic protein (MBP). Iba1 was used to visualise microglia along with a surrogate marker of phagocytosis, CD68, and one of the key components of myelin, MBP, to investigate microglial phagocytosis of myelin following injury and between biological sexes. Microglia colocalisation with CD68 and MBP was used in this study as a measure of microglial phagocytosis of myelin. Microglial phagocytosis of myelin was examined in the corpus callosum (CC; Figure 6) and primary sensory barrel field (S1BF; Figure 7), where CD68 staining was most prevalent, and was reported in this study as the proportion of microglia colocalised with CD68 and MBP +/- 95% CI. Regional differences in the proportion of CD68-positive/MBP-positive (CD68+/MBP+) microglia were observed across time post-injury and between biological sexes (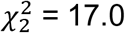; *p* < 0.0001) (see Supplementary Table 1 for estimated counts and their 95% Cis; depicted in Figure 6 and 7). In the CC, there was a significant reduction in the proportion of CD68+/MBP+ microglia at 3 days post-injury (DPI) compared with naïve controls in both males (*p* = 0.0062; see Table 3; Figure 6) and females (*p* = 0.0265; see Table 3; Figure 6). There was a significant increase in the proportion of CD68+/MBP+ microglia in the CC of male mice at 7 days post-injury (DPI) compared with sex-matched naïve controls (*p* = 0.0154; see Table 3; Figure 6) and at 3 days post-injury (*p* < 0.0001; see Table 3; Figure 6) but not in females. The proportion of CD68+/MBP+ microglia in the CC of male mice was also significantly greater than in females at 7 days post-injury (*p* = 0.0005; see Table 4; Figure 6). In the S1BF there was a significant decrease in the proportion of CD68+/MBP+ microglia at 3 days post-injury in males (*p* = 0.0260; see Table 3; Figure 7) whilst there was a significant increase in females (*p* = 0.0044; see Table 3; Figure 7) compared with sex-matched naïve controls. The proportion of CD68+/MBP+ microglia was also significantly greater in the S1BF of females compared with males at 3 days post-injury (*p* < 0.0001; see Table 4; Figure 7). At 7 days post-injury, there was a significant increase in the proportion of CD68+/MBP+ microglia in the S1BF of males compared with sex-matched naïve controls (*p* = 0.0007; see Table 3; Figure 7) and at 3 days post-injury (*p* < 0.0001; see Table 3; Figure 7). However, there was a significant reduction in the proportion of CD68+/MBP+ microglia in the S1BF of females at 7 days post-injury compared with sex-matched naïve controls (*p* = 0.0132; see Table 3; Figure 7) and at 3 days post-injury (*p* < 0.0001; see Table 3; Figure 7). The proportion of CD68+/MBP+ microglia was also significantly greater in the S1BF of males compared with female mice at 7 days post-injury (*p* = 0.0001; see Table 4; Figure 7). These results suggest that male mice exhibit a reduction in microglial phagocytosis of myelin at 3 days post-injury in both the CC and S1BF that becomes increased at 7 days post-injury above naïve levels. Female mice also experience a reduction in microglial phagocytosis of myelin at 3 days post-injury in the CC that returns to naïve levels by 7 days post-injury. However, in the S1BF, female mice exhibit an increase in microglial phagocytosis at 3 days post-injury which becomes reduced below naïve levels at 7 days post-injury.

**Figure 6:**
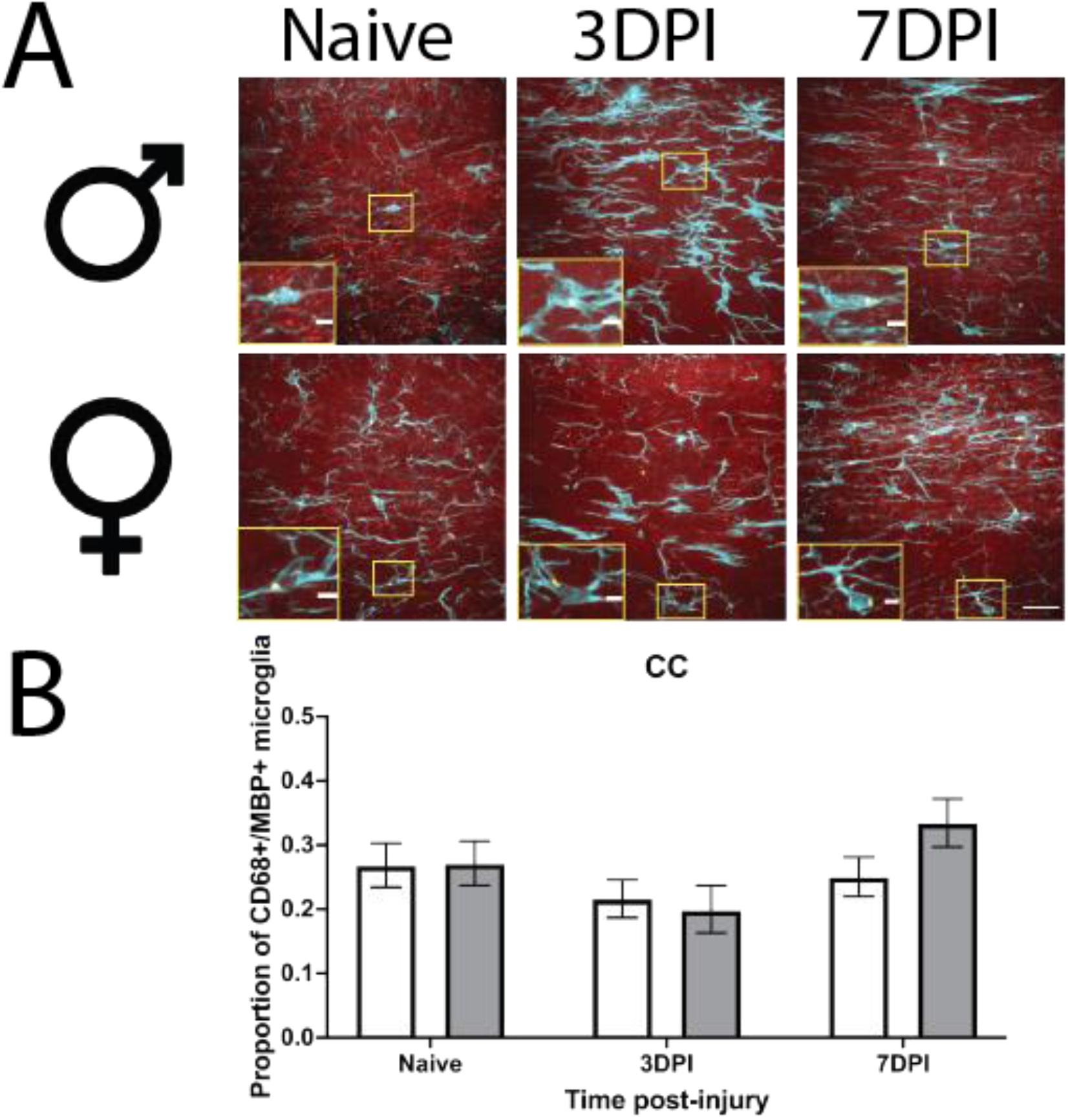
Example images of microglia colocalised with CD68 and MBP in the corpus callosum (CC) of male and female mice in a naïve state or at 3- or 7-days post-injury (DPI; **A**; scale bar = 25 μm, inset = 5 μm). Microglial colocalisation with CD68 and MBP was quantified in the CC of both males (grey) and females (white) and expressed as the proportion of CD68+/MBP+ microglia (mean ±95%CI; **B**). There was a significant reduction in the proportion of CD68+/MBP+ microglia at 3 days post-injury in both males and females compared with sex-matched naïve controls (see Table 3). At 7 days post-injury, there was a significant increase in the proportion of CD68+/MBP+ microglia in males compared with sex-matched naïve controls and at 3 days post-injury, which did not occur in females (see Table 3). The proportion of CD68+/MBP+ microglia was also significantly greater in males compared with females at 7 days post-injury (see Table 4).

**Figure 7:**
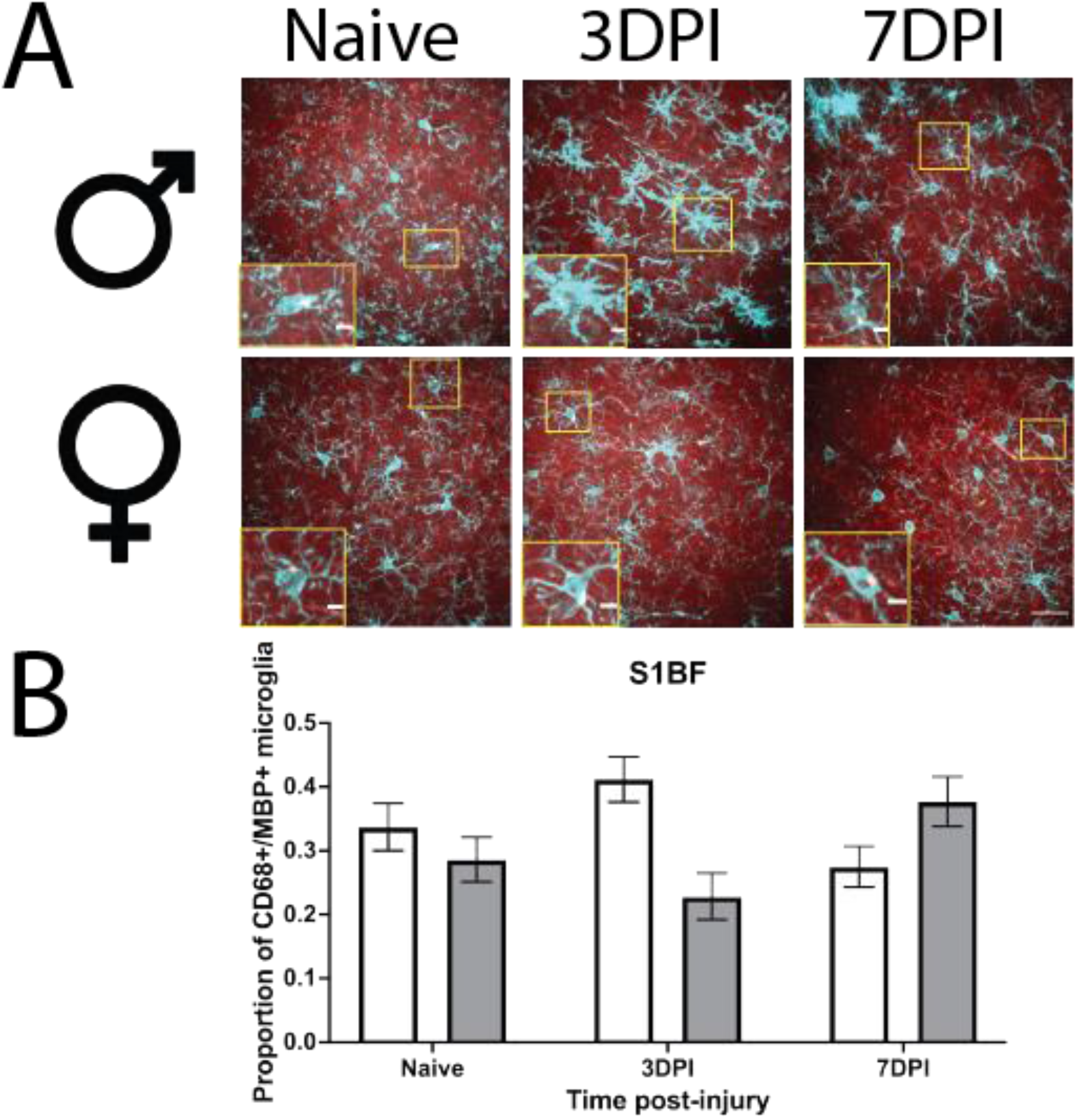
Example images of microglia colocalised with CD68 and MBP in the primary sensory barrel field (S1BF) of male and female mice in a naïve state or at 3- or 7-days post-injury (DPI; **A**; scale bar = 25 μm, inset = 5 μm). Microglial colocalisation with CD68 and MBP was quantified in the S1BF of both males (grey) and females (white) and expressed as the proportion of CD68+/MBP+ microglia (mean ±95%CI; **B**). There was a significant reduction in the proportion of CD68+/MBP+ microglia at 3 days post-injury in males compared with sex-matched naïve controls (see Table 3). Whilst females experienced a significant increase in the proportion of CD68+/MBP+ microglia at 3 days post-injury compared with sex-matched naive controls (see Table 3). The proportion of CD68+/MBP+ microglia was also significantly greater in females compared with males at 3 days post-injury (see Table 4). At 7 days post-injury, there was a significant increase in the proportion of CD68+/MBP+ microglia in males compared with sex-matched naïve controls and at 3 days post-injury (see Table 3). However, females exhibited a significant reduction in the proportion of CD68+/MBP+ microglia at 7 days post-injury compared with sex-matched naïve controls and at 3 days post-injury (see Table 3). The proportion of CD68+/MBP+ microglia was also significantly greater in males compared with females at 7 days post-injury (see Table 4).

**Table 3:** Colocalised microglia post-hoc contrasts across time post-injury, separated by region and biological sex, using Tukey’s procedure. Bold means statistically significant p < 0.05.

| Contrast:<br>Injury (Tukey) | Region | Sex | Co-efficient<br>estimate | Lower 95% CI | Upper 95% CI | p-values |
| --- | --- | --- | --- | --- | --- | --- |
| N vs 3DPI | CC | F | 0.2825 | 0.0330 | 0.5321 | **0.0265** |
| N vs 7DPI | CC | F | 0.0915 | -0.1471 | 0.3302 | 0.4522 |
| 3DPI vs 7DPI | CC | F | -0.1910 | -0.4298 | -0.0478 | 0.1169 |
| N vs 3DPI | S1BF | F | -0.3231 | -0.5456 | -0.1007 | **0.0044** |
| N vs 7DPI | S1BF | F | 0.2937 | 0.0614 | 0.5260 | **0.0132** |
| 3DPI vs 7DPI | S1BF | F | 0.6169 | 0.3967 | 0.8371 | **<0.0001** |
| N vs 3DPI | CC | M | 0.4098 | 0.1166 | 0.7031 | **0.0062** |
| N vs 7DPI | CC | M | -0.3015 | -0.5454 | -0.0576 | **0.0154** |
| 3DPI vs 7DPI | CC | M | -0.7113 | -0.9991 | -0.4236 | **<0.0001** |
| N vs 3DPI | S1BF | M | 0.3073 | 0.0368 | 0.5778 | **0.0260** |
| N vs 7DPI | S1BF | M | -0.4148 | -0.6554 | -0.1742 | **0.0007** |
| 3DPI vs 7DPI | S1BF | M | -0.7221 | -0.9896 | -0.4546 | **<0.0001** |

**Table 4:**
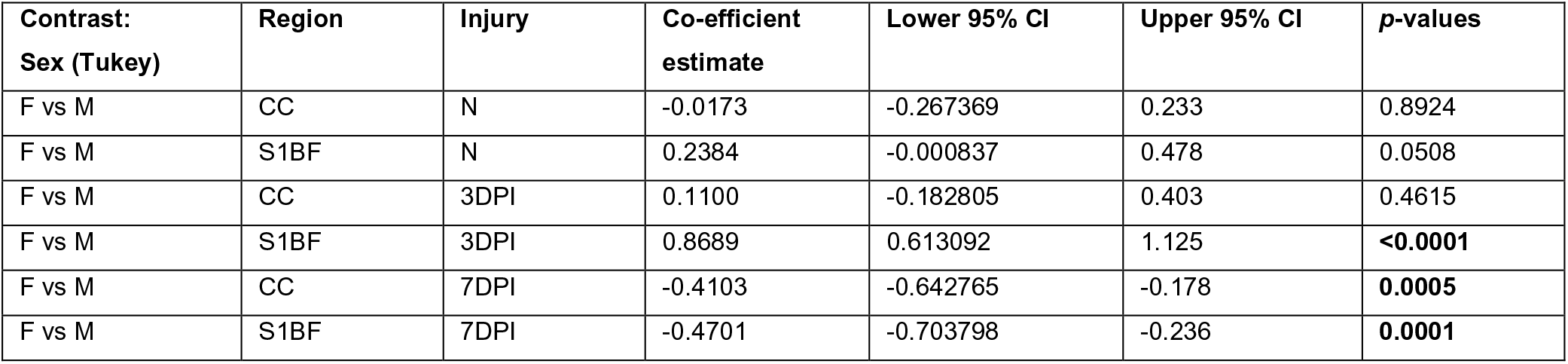
Colocalised microglia post-hoc contrasts between males and females, separated by region and injury status, using Tukey’s procedure. Bold means statistically significant p < 0.05.

| Contrast:<br>Sex (Tukey) | Region | Injury | Co-efficient<br>estimate | Lower 95% CI | Upper 95% CI | p-values |
| --- | --- | --- | --- | --- | --- | --- |
| F vs M | CC | N | -0.0173 | -0.267369 | 0.233 | 0.8924 |
| F vs M | S1BF | N | 0.2384 | -0.000837 | 0.478 | 0.0508 |
| F vs M | CC | 3DPI | 0.1100 | -0.182805 | 0.403 | 0.4615 |
| F vs M | S1BF | 3DPI | 0.8689 | 0.613092 | 1.125 | **<0.0001** |
| F vs M | CC | 7DPI | -0.4103 | -0.642765 | -0.178 | **0.0005** |
| F vs M | S1BF | 7DPI | -0.4701 | -0.703798 | -0.236 | **0.0001** |

### Microglia isolated from mice at 28 days post-injury had a reduction in pro-inflammatory proteins and an increase in anti-inflammatory proteins compared with naive controls

The first SPC scores (Figure 2) were significantly decreased in microglia isolated from mice at 28 days following mFPI compared with naïve levels (see Table 1; Figure 3). Using STRING gene ontology (GO) analysis, SPC1 protein communities were related to metabolic processes (Figure 8A). STRING Wikipathway analysis revealed that SPC1 protein communities were involved with pro-inflammation via the TNF-α NF-κβ signalling pathway (“tumour necrosis factor alpha (TNF-α) nuclear factor kappa beta (NF-κβ) signalling pathway:WP246”) along with protein (“proteasome degradation:WP519”) and mRNA metabolism (“mRNA processing:WP310”; Figure 8B). SPC1 protein communities were also involved with proliferation (“IL-2 signalling pathway:WP450”, “IL-3 signalling pathway:WP373” and “EGFR1 signalling pathway:WP572”) and purine metabolism (“purine metabolism:WP2185”) that was determined via Wikipathway analysis (Figure 8B). ATP is a type of purine that is released by damaged neurons to trigger the migration of microglia to the injury site and the metabolism of ATP by microglia promotes phagocytic and pro-inflammatory functions (Jassam et al. 2017). Furthermore, SPC1 protein communities were involved with amino acid metabolism (“amino acid metabolism:WP662”), fatty acid oxidation (“fatty acid β-oxidation:WP1269”, “mitochondrial fatty acid β-oxidation:WP401”) and oxidative phosphorylation (“oxidative phosphorylation:WP1248”, “electron transport chain:WP295”, “TCA cycle:WP454”). These results suggest that amino acid metabolism, fatty acid oxidation and oxidative phosphorylation are associated with microglial pro-inflammation which is in line with previous literature that amino acid metabolism is utilised during pro-inflammatory functions (Geric et al. 2019). However, fatty acid oxidation is typically used by homeostatic and anti-inflammatory microglia to fuel house-keeping functions and tissue remodelling (reviewed in Yang et al. 2021). Studies in macrophages have shown that fatty acid oxidation was important for the release of pro-inflammatory cytokines such as IL-1*β* and IL-18 (Moon et al. 2016). These findings could have implications for microglial metabolic programming in relation to pro-inflammatory functions. As SPC1 scores were significantly decreased at 28 days post-injury from naïve levels, our findings indicate a reduction in microglial pro-inflammation at 28 days post-injury.

**Figure 8:**
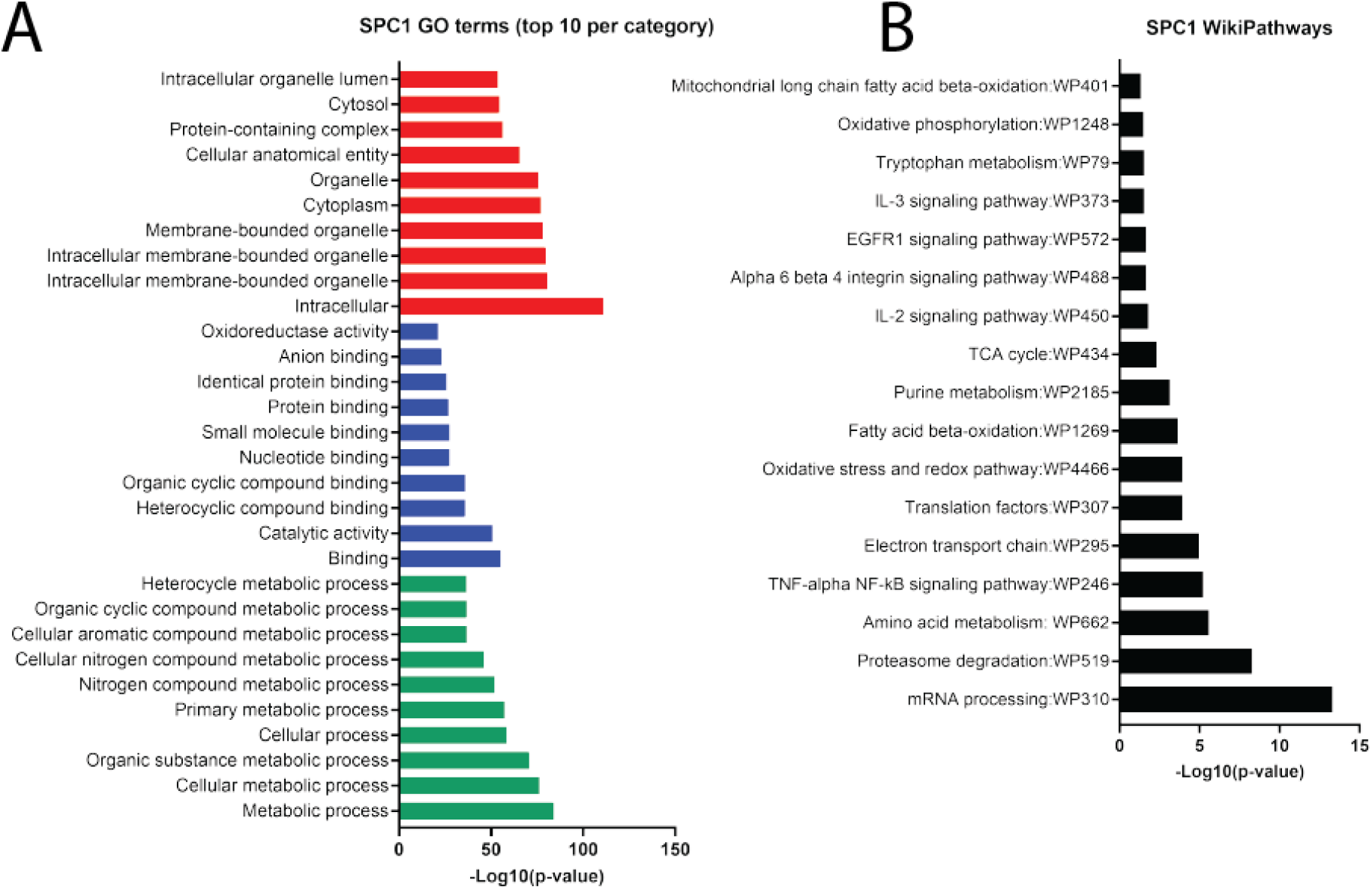
STRING gene ontology (GO; A) and WikiPathway (B) analysis of sparse principal component 1 (SPC1) protein communities. GO analysis includes the top 10 GO terms per category; biological process (green), molecular function (blue) and cellular component (red). Made with GraphPad Prism, version 8.

However, the second SPC scores (Figure 2) were significantly increased in microglia isolated from mice at 28 days following mFPI compared with naïve levels (see Table 1; Figure 3). STRING gene ontology (GO) analysis determined that SPC2 protein communities were related to gene expression and mRNA processing (Figure 9A). STRING Wikipathway analysis also revealed that SPC2 protein communities were involved with pluripotency (“PluriNetWork: mechanisms associated with pluripotency:WP1763”) and protein (“cytoplasmic ribosomal proteins:WP163”) and mRNA metabolism (“mRNA processing:WP310”) (Figure 9B). SPC2 protein communities were also related to TNF-α NF-κβ signalling (“TNF-α NF-κβ signalling pathway:WP246”) which encompassed the A-kinase anchoring protein 8 (Akap8; see Table 5). Akap8, also known as Akap95, is a protein that modulates toll-like receptor signalling and inhibits TNF-α NF-κβ mediated pro-inflammation (Wall et al. 2009). SPC2 protein communities related to the TNF-α NF-κβ signalling were also distinct from SPC1 protein communities, indicating that SPC1 and SPC2 protein communities are involved with different TNF-α NF-κβ signalling cascades (see Table 5; Figure 10). Therefore, microglial protein communities increased at 28 days post-injury were involved with anti-inflammatory functions via TNF-α NF-κβ signalling. SPC2 protein communities were also related to Wikipathways involved with oxidative phosphorylation (“electron transport chain:WP295”; Figure 9B). These results indicate that oxidative phosphorylation is associated with microglial anti-inflammatory function which is in line with previous literature (Holland et al. 2018; reviewed in Yang et al. 2021). As SPC2 scores were significantly increased at 28 days post-injury compared with naïve levels, these results suggest that anti-inflammatory proteins were increased in microglia at 28 days post-injury compared with naïves. Therefore, this data indicates that microglial pro-inflammation was reduced at 28 days post-injury whilst anti-inflammatory function was increased.

**Figure 9:**
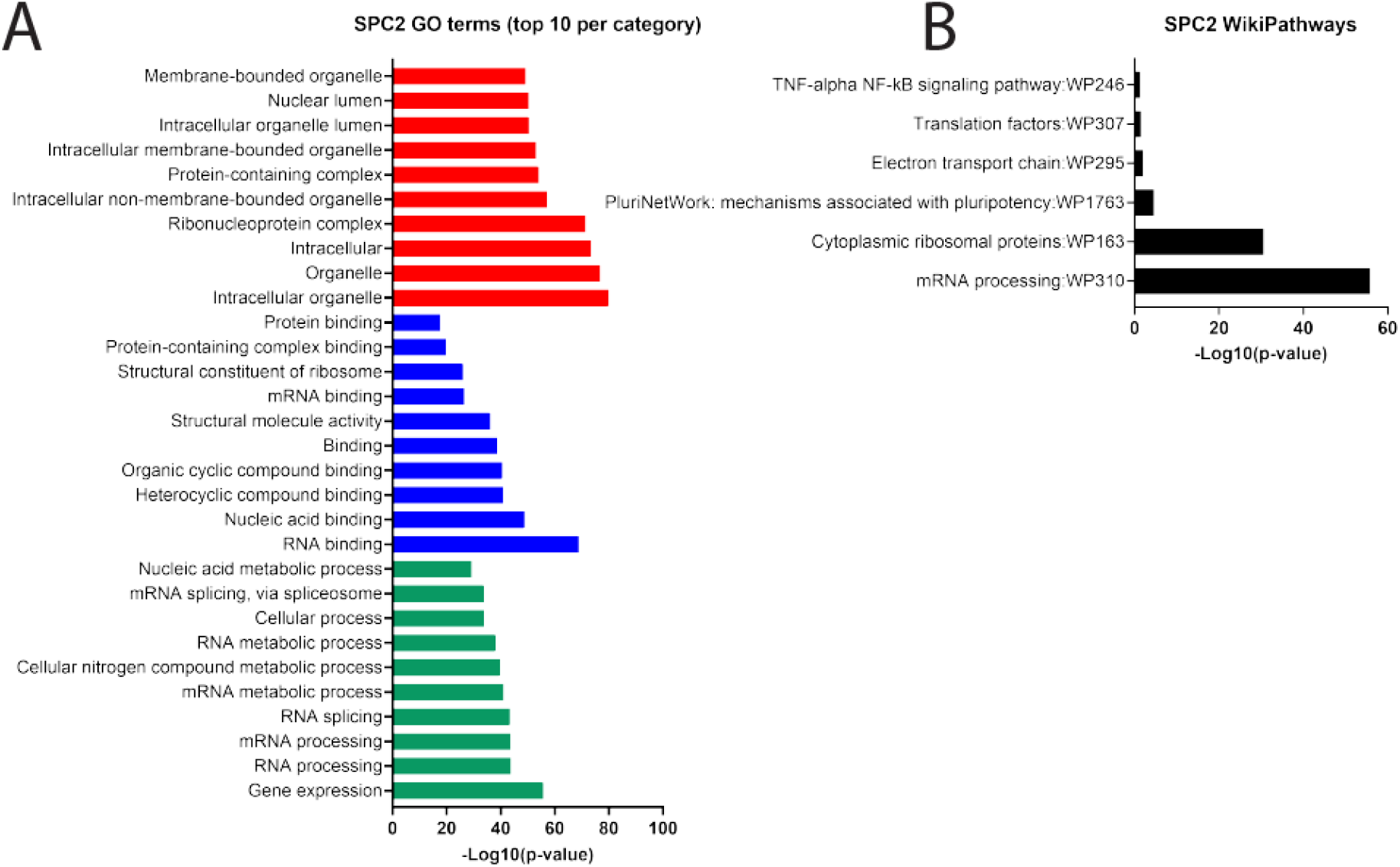
STRING gene ontology (GO; A) and WikiPathway (B) analysis of second sparse principal component 2 (SPC2) protein communities. GO analysis includes the top 10 GO terms per category; biological process (green), molecular function (blue) and cellular component (red). Made with GraphPad Prism, version 8.

**Figure 10:**
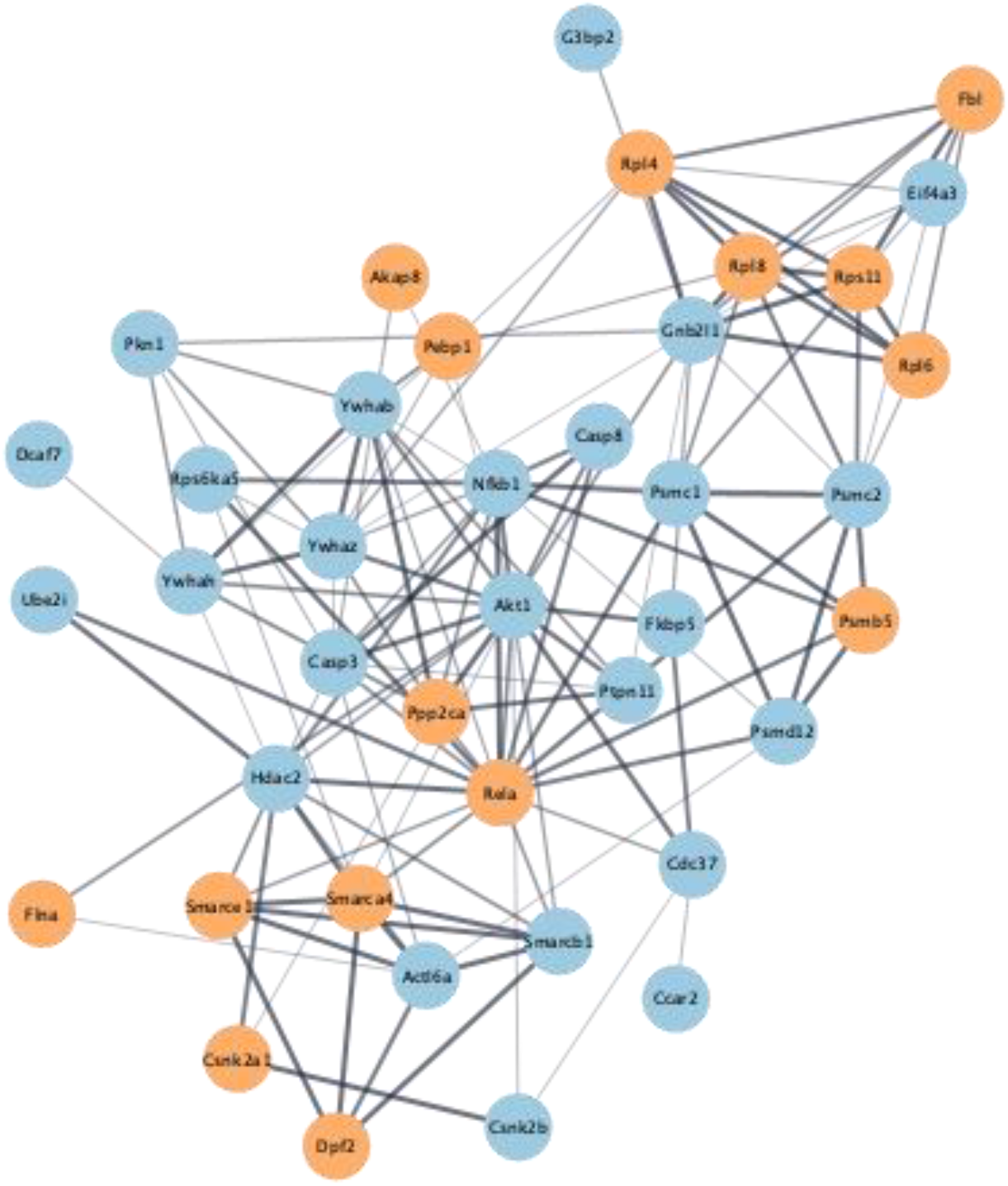
STRING protein network of the first (blue) and second (orange) sparse principal component 1 (SPC1 - blue) and SPC2 (orange) protein communities that were related to TNF-α NF-κβ pro- and anti-inflammation (“TNF-α NF-κβ signalling pathway:WP246”), respectively, determined via STRING Wikipathway analysis.

**Table 5:**
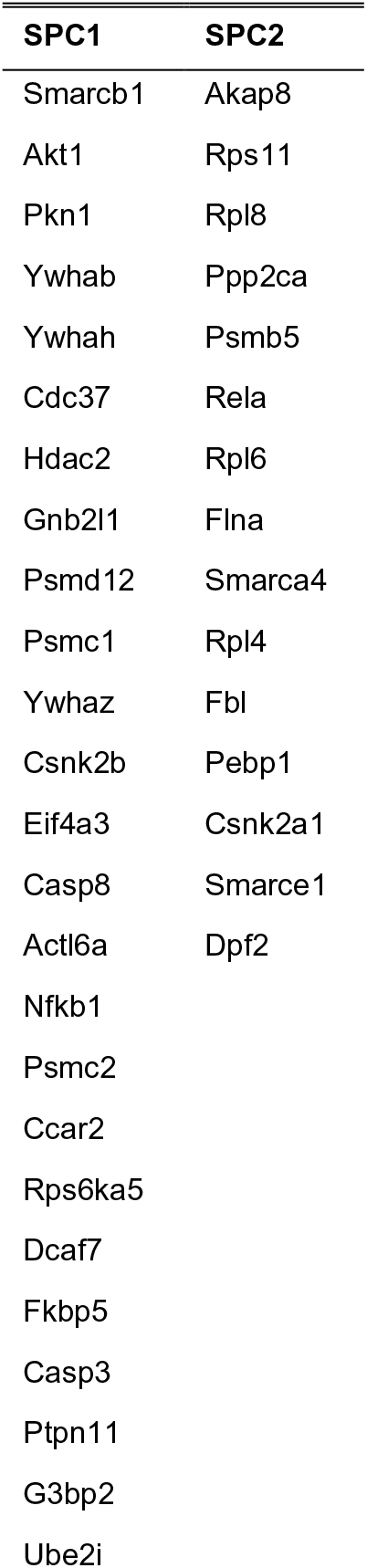
Sparse principal component 1 (SPC1) and SPC2 protein communities that were related to TNF-α NF-κβ pro- and anti-inflammation (“TNF-α NF-κβ signalling pathway:WP246”), respectively, identified via STRING WikiPathway analysis.

### Microglia isolated from male mice had an increase in protein communities related to pro-inflammation and phagocytosis as well as estrogen and insulin signalling compared with females

The fourth SPC scores (Figure 2) were significantly increased in microglia isolated from male mice compared with females, regardless of injury (see Table 1). Using STRING gene ontology (GO) analysis, SPC4 protein communities were related to cellular and metabolic processes (Figure 11A). SPC4 protein communities were identified via Wikipathways to be involved with pro-inflammation (“TNF-α NF-κβ signalling pathway:WP246”, “IL-6 signalling pathway:WP387”), proliferation (“IL-2 signalling pathway:WP450”, “IL-3 signalling pathway:WP373”, “IL-5 signalling pathway:WP151”, “IL-7 signalling pathway:WP297”, “IL-9 signalling pathway:WP10”, “EGFR1 signalling pathway:WP572”), cell migration (“chemokines signalling pathway:WP2292”, “purine metabolism:WP2185”) and phagocytosis (“regulation of actin cytoskeleton:WP523”, “microglia pathogen phagocytosis pathway:WP3626”; Figure 11B). Protein communities were also related to tyrosine kinase-binding protein (Tyrobp) signalling (“Tyrobp causal network in microglia:WP3625”) identified via Wikipathway analysis (Figure 11B). Tyrobp is also known as DAP12 and is reported to be involved with proliferative and phagocytic functions in addition to being linked to the pathogenesis of Alzheimer’s disease (AD) (Haure-Mirande et al. 2019; Yao et al. 2019). Additionally, serotonin signalling (“serotonin and anxiety:WP2141”) was related to SPC4 protein communities which has been previously reported to enhance microglial migration to ATP, but, conversely, attenuates phagocytosis (Krabbe et al. 2012; reviewed in D’Andrea, Béchade, and Maroteaux 2020). SPC4 protein communities were also related to oxidative stress (“oxidative stress and redox pathway:WP4466) which has been extensively reported to be triggered by pro-inflammation (reviewed in Lushchak et al. 2021). Therefore, microglia from males exhibited an increase in pro-inflammatory and phagocytic proteins compared with females which involved Tyrobp and serotonin signalling in addition to oxidative stress. Furthermore, amino acid metabolism (“amino acid metabolism:WP662”) and glycolysis (“pentose phosphate pathway:WP63) were related to SPC4 protein communities. These results suggest that amino acid metabolism and glycolysis are associated with pro-inflammation and phagocytosis which is in line with previous literature (Wang et al. 2019; Geric et al. 2019; reviewed in Lauro and Limatola 2020). Insulin (“insulin signalling:WP65”) and estrogen signalling (“estrogen signalling:WP1244”) were also related to SPC4 protein communities that was determined via Wikipathway analysis (Figure 11B). Notably, the protein communities related to pro-inflammation (“TNF-α NF-κβ signalling pathway:WP246”) and insulin (“insulin signalling:WP65”) and estrogen signalling (“estrogen signalling:WP1244”) all contained the Akt1 protein, suggesting that Akt1 may be involved in microglial biological sex differences of these pathways (see Table 6, Figure 12). As the fourth SPC scores were increased in microglia isolated from males compared with females, this data indicates microglial pro-inflammation and phagocytosis as well as estrogen and insulin signalling was increased in male mice compared with females.

**Figure 11:**
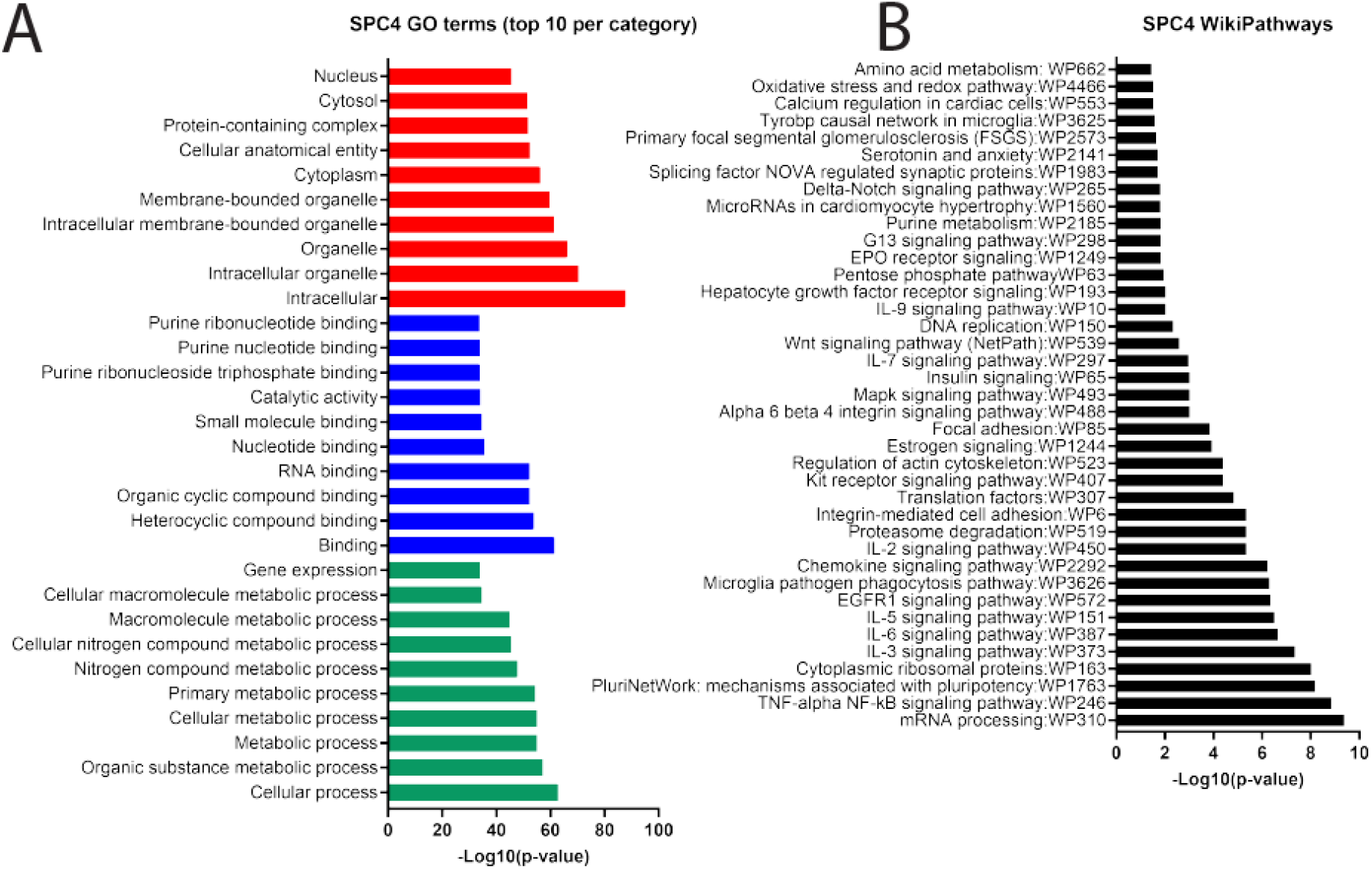
STRING gene ontology (GO; A) and WikiPathway (B) analysis of sparse principal component 4 (SPC4) protein communities. GO analysis includes the top 10 GO terms per category; biological process (green), molecular function (blue) and cellular component (red). Made with GraphPad Prism, version 8.

**Figure 12:**
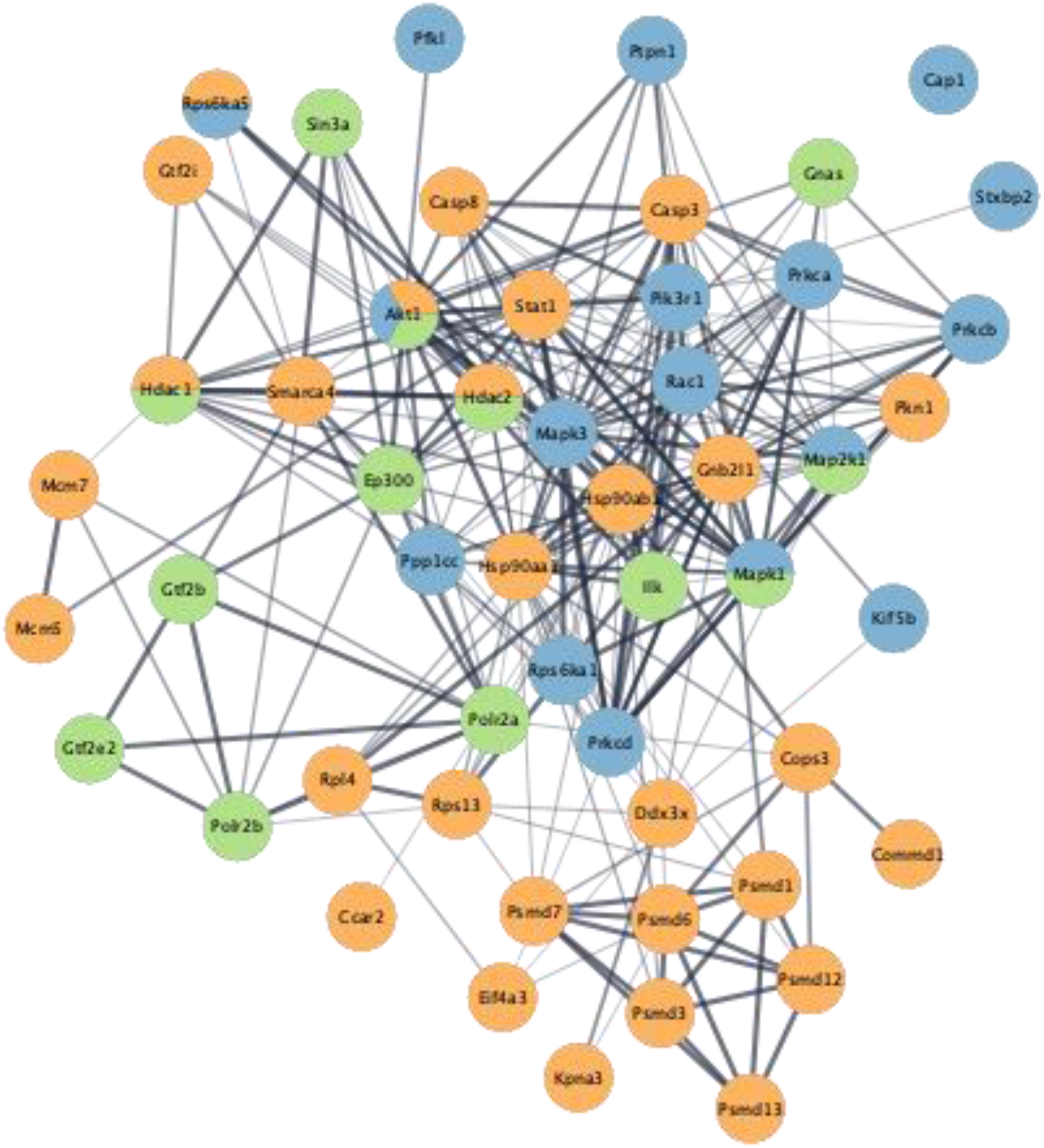
STRING protein network of sparse principal component 4 (SPC4) protein communities related to pro-inflammation (“TNF-α NF-κβ signalling pathwayWP426” - orange) as well as insulin (“insulin signalling:WP65” - blue) and estrogen signalling (“estrogen signalling:WP1244” - green) and identified via STRING Wikipathway analysis.

**Table 6:**
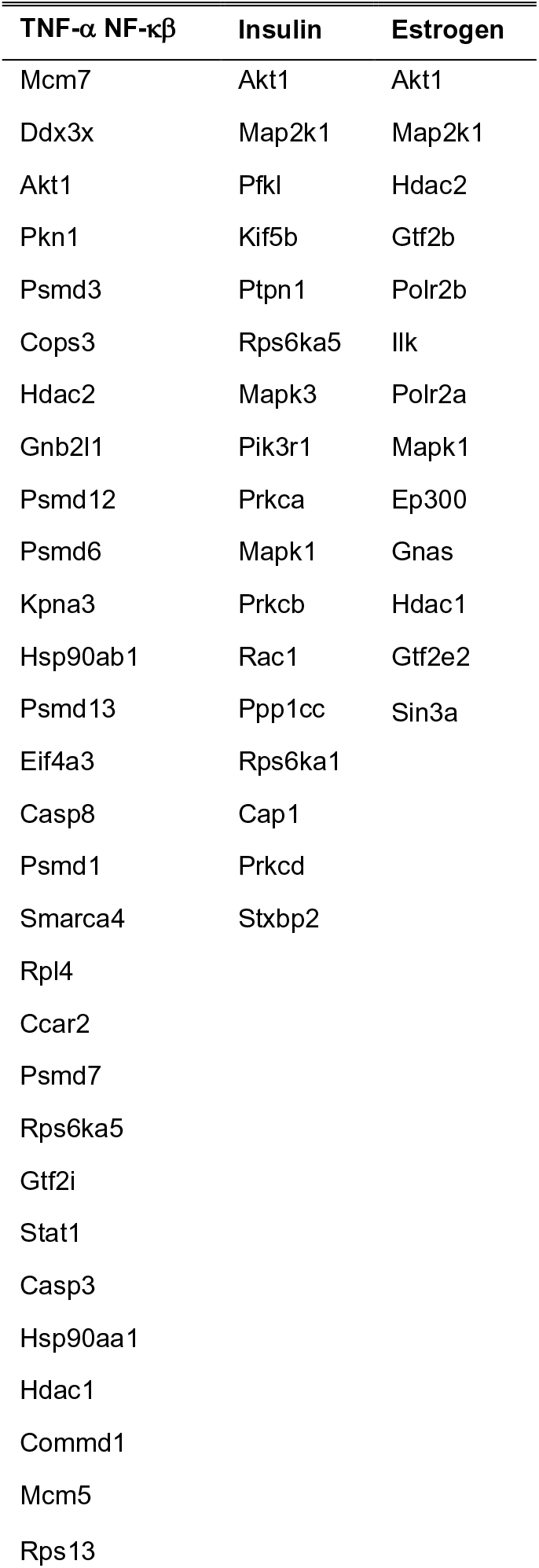
Sparse principal component 4 (SPC4) protein communities related to TNF-α NF-κβ pro-inflammation (“TNF-α NF-κβ signalling pathway:WP246”) as well as insulin (“insulin signalling:WP65”) and estrogen signalling (“estrogen signalling:WP1244”) identified via STRING WikiPathway analysis.

## Discussion

It is currently unclear why certain individuals are susceptible to developing long-term symptoms after a TBI but it may be related to persistent neuroinflammation mediated by microglia. The microglial response is typically thought to peak in the days to weeks following TBI. This response is encompassed by morphological changes along with pro- and anti-inflammation as well as phagocytosis which is beneficial for tissue repair (Rowland et al. 2020; Smith et al. 2013; Loane et al. 2014; Cao et al. 2012). However, there is increasing literature that has demonstrated microglial pro-inflammation and phagocytosis can be evident for years post-injury and is associated with worsened neuropathological and functional outcomes (Loane et al. 2014; Ritzel et al. 2020; Boone et al. 2019; Johnson et al. 2013). Microglia have also been reported to exhibit biological sex differences in the healthy brain as well as after a TBI which may impact TBI recovery (Doran et al. 2019; Villapol, Loane, and Burns 2017; Yanguas-Cas·s 2020; Villa et al. 2018; Guneykaya et al. 2018). Therefore, this study sort to identify a distinct pattern of microglial functions across time after a TBI and whether this differs between biological sexes. To do this, we examined the microglial proteome in the days (3- and 7-days) to 1 month (28 days) following a single diffuse TBI (mFPI) in male and female mice. Our core findings relate to temporal changes in microglial proteins associated with phagocytosis, inflammation and metabolism/energy homeostasis across time post-injury.

Overall, the results from this study showed a reduction in microglial phagocytosis of neuronal elements at 3- and 7-days post-injury compared with naïve, with a shift to anti-inflammatory functions by 28 days in both biological sexes. This is in contrast to previous studies that showed elevated microglial pro-inflammation and phagocytosis within days up to years following TBI in humans and rodents using immunohistochemistry, quantitative real-time (qRT)-PCR, autoradiography or flow cytometry (Kelley, Lifshitz, and Povlishock 2007; Ritzel et al. 2020; Johnson et al. 2013; Cao et al. 2012; Madathil et al. 2018; Witcher et al. 2021). No increase in microglial proteins related to pro-inflammation and phagocytosis across time post-injury may be due to the global analysis conducted in this study. However, this is one of the first studies to investigate global microglial protein changes across time post-injury in both biological sexes. Others have either concentrated on specific regions of the brain, including the cortex, hippocampus and thalamus, or only included male rodents (Kelley, Lifshitz, and Povlishock 2007; Ritzel et al. 2020; Johnson et al. 2013; Cao et al. 2012; Madathil et al. 2018; Witcher et al. 2021). One study which performed a global analysis of temporal changes in microglial mRNA at 2-, 14- and 60-days after a focal TBI (CCI) in male mice via qPCR reported a reduction in mRNA related to microglial sensing abilities and homeostatic functions at 2 days post-injury (Izzy et al. 2019). How this translates to protein levels remains unknown. However, a reduction in microglial sensing capabilities in the days following TBI could impact upon microglial phagocytic functions and, therefore, explain why we observed a reduction in phagocytic proteins. Furthermore, as microglia phagocytose neuronal elements, including myelin and synapses, under homeostatic conditions in order to refine neural circuitry (reviewed in Galloway et al. 2019; Santos and Fields 2021), a reduction in homeostatic functions observed at 2 days post-injury in the previous study could result in a reduction in phagocytosis which we observed at 3- and 7-days post-injury. Previous literature has also reported a mixed mode of microglial inflammation after a TBI (Kumar et al. 2016; Morganti, Riparip, and Rosi 2016). Our data, taken in conjunction with Izzy et al. 2019, indicates peaks and troughs in microglial pro- and anti-inflammation across time after a TBI. The peaks could shift to a pro-inflammatory profile in the months to years post-TBI. However, we only examined the microglial proteome up to 1 month post-injury, therefore, we were unable to determine whether there were changes in the microglial proteome months to years post-injury and whether these differed between biological sexes.

A number of studies have reported that males exhibit a greater inflammatory and phagocytic response in the short-term following TBI (Villapol, Loane, and Burns 2017; Bromberg et al. 2020). However, we found no biological sex differences in the microglial proteome across time post-injury suggesting that the microglial response is similar between males and females up to 1 month post-injury. This is in line with previous literature that did not observe any significant differences in the expression of pro- or anti-inflammatory cytokines, production of reactive oxygen species (ROS) or microglial phagocytic activity following a CCI between male and female mice (Doran et al. 2019). However, Doran et al., 2019 did identify an influx of peripheral myeloid cells and microglial proliferation at 1- and 3-days post-injury in male mice, determined using flow cytometry, which did not occur in females. Therefore, the infiltration of peripheral myeloid cells in males but not females may be contributing to the biological sex differences in neuroinflammation reported in other studies that has been previously thought to be mediated by microglia. As we only examined the microglial response following TBI, it is unclear whether peripheral macrophages were contributing to the neuroinflammatory response in the current study and if this differs between biological sex differences. Although no injury specific differences were present in our study between biological sexes, we did observe an increase in pro-inflammatory and phagocytic proteins in microglia isolated from male mice compared with females. This was accompanied by an increase in microglial proteins related to insulin and estrogen signalling in male mice compared with females. Notably, the Akt1 protein was increased in microglia isolated from males compared with females which was involved with pro-inflammation as well as insulin and estrogen signalling pathways. Protein kinase B serine/threonine kinase 1 (Akt1) is the primary isoform of Akt expressed in microglia that has been shown, *in vitro* and *in vivo*, to become activated via toll-like receptor 4 (TLR4) and promote the production of TNF-α via the phosphorylation of NF-κβ (Huang et al. 2016; Lee et al. 2006; Xu et al. 2020; Saponaro et al. 2012; Levenga et al. 2021) Insulin has also been shown, *in vivo*, to increase microglial expression of pro-inflammatory mediators (COX2/IL-1β) via Akt signalling cascades in the hippocampus of young male rats, but not aged animals (Haas et al. 2020). *In vitro* studies have also shown that insulin induces the production of pro-inflammatory cytokines by microglia and increases phagocytic activity (Brabazon et al. 2018; Spielman et al. 2015). Therefore, these data suggest that insulin may promote a pro-inflammatory and phagocytic profile of microglia via Akt1 intracellular signalling in males, but not females.

Estrogen is known as one of the primary female sex hormones, however, males also produce estrogen at a lower levels to that of females (reviewed in Gillies and McArthur 2010). The role of estrogen in the brain has been extensively reported to exhibit anti-inflammatory properties and promote cell survival and neuroplasticity under homeostatic and pathological conditions, such as after a TBI (Wang et al. 2021; Honig et al. 2021; reviewed in Kövesdi, Szabó-Meleg, and Abrahám 2020). However, estrogen synthesised from testosterone during development has been shown to induce microglial morphological changes and the production of pro-inflammatory cytokines, particularly prostaglandin E_2_ (PGE_2_), which is integral in the masculinisation of the male brain (Lenz et al. 2013). A reduction in testosterone and an increase in estrogen production in young and older men has also been correlated with increased systemic pro-inflammatory cytokines (Bobjer et al. 2013; Maggio et al. 2009; Barud et al. 2010). Therefore, estrogen synthesised from testosterone may promote pro-inflammatory functionality of microglia in the adult male brain. However, these pathways are extremely complex and our current study design cannot determine what is driving the changes in each pathway. Further studies are required to examine the effects of insulin and estrogen signalling upon microglial functionality in the healthy brain of both males and females as well as after a TBI, especially given endocrine dysfunction post-TBI (reviewed in Li and Sirko, 2018).

One of the limitations of this study is that we conducted a global analysis of microglial proteins from the whole brain, thus, we are a capturing an average of microglial function across the entire brain. Microglia are known to exhibit heterogeneity between brain regions including density, morphology and expression of proteins that are evident in the healthy brain and in response to stimuli, such as a TBI (Stratoulias et al. 2019; Tan, Yuan, and Tian 2020; Böttcher et al. 2019). Therefore, it may only be microglia in particular brain regions that are participating in pro-inflammatory and phagocytic functions following TBI. To test this, we performed immunohistochemistry to examine the spatial relationship in microglial phagocytosis of myelin across time post-injury and between biological sexes. We observed regional differences in the proportion of microglia colocalised with a surrogate marker of phagocytosis (CD68) and one of the key constituents of myelin (MBP) in females but not males. Males exhibited a reduction in microglial colocalisation with CD68 and MBP at 3 days post-injury compared to naïve which became increased above naïve levels at 7 days in both the CC and S1BF. Females also exhibited a reduction in microglial colocalisation at 3 days post-injury in the CC, but the opposite trend was observed in the S1BF where microglial colocalisation was increased above naïve levels at 3 days and became reduced at 7 days post-injury. These findings suggest that subtle regional changes in microglial phagocytosis of neuronal elements occur across time post-injury and between biological sexes that were not able to be detected using a global whole brain analysis. An increase in microglial pro-inflammation after a TBI may also be dependent upon brain region but we did not examine this in the current study. Further research is required to better understand how microglial heterogeneity contributes to the microglial response under pathological conditions. Another limitation of this work is that we did not examine whether TBI induced neuropathology or behavioural impairments was correlated to proteome changes. Previous studies using male rodents have shown that a single diffuse TBI can induce neuron loss in the sensory cortex in addition to motor and neurological deficits in the days post-injury (Lifshitz and Lisembee 2012; Rowe et al. 2019). Our results do show a reduction in microglial phagocytosis of neuronal elements at 3- and 7-days post-injury in both male and female mice. Myelin damage in the corpus callosum has been associated with cognitive dysfunction days to months after a TBI in male mice (Nonaka et al. 2021; Taib et al. 2017). However, we were not able to determine whether changes in microglial phagocytosis after a TBI contributes to neuropathology or myelin damage post-injury and whether this differs between biological sexes.

In conclusion, the findings from this study demonstrated that microglia undergo temporal changes in phagocytic as well as pro- and anti-inflammatory proteins following a diffuse TBI in male and female mice. Temporal changes in the microglial proteome across time post-injury may impact TBI recovery, however, additional studies are required to determine whether these changes correlate with neuropathological and functional outcomes. Despite the changes microglial proteins across time post-injury being similar between biological sexes, we observed an increase in pro-inflammatory and phagocytic proteins in males compared with females regardless of injury. This pro-inflammatory and phagocytic profile of microglia in males was accompanied by an increase in insulin and estrogen signalling proteins compared with females. The increased insulin and estrogen signalling observed in males could be driving microglial pro-inflammation and phagocytosis. However, further research is required to understand the roles of insulin and estrogen upon microglial function between biological sexes and how this impacts the microglial response following TBI. Even though the microglial response across time post-injury was similar between males and females, biological sex differences in basal microglial functionality may impact the effectiveness of treatment strategies targeting the microglial response post-injury.

## Acknowledgements

The authors would like to thank Dr. Terry L. Pinfold and Dr. William R. Bennett for their technical assistance and Dr. Dino Premilovac for his assistance with data interpretation.

