## Supplemental Table 1 for "Temporal changes in the microglial proteome of male and female mice after a diffuse brain injury using label-free quantitative proteomics"

### Supplementary material

#### 1. Supplementary tables

**Supplementary Table 1:** *Colocalised microglia were counted in the corpus callosum (CC) and primary sensory barrel field (S1BF).*

| Injury | Region | Sex | Probability | Lower 95% CI | Upper 95% CI |
|---|---|---|---|---|---|
| N | CC | F | 0.267 | 0.234 | 0.303 |
| N | S1BF | F | 0.336 | 0.300 | 0.374 |
| 3 | CC | F | 0.215 | 0.187 | 0.246 |
| 3 | S1BF | F | 0.411 | 0.376 | 0.447 |
| 7 | CC | F | 0.249 | 0.220 | 0.281 |
| 7 | S1BF | F | 0.274 | 0.243 | 0.307 |
| N | CC | M | 0.270 | 0.237 | 0.306 |
| N | S1BF | M | 0.285 | 0.251 | 0.321 |
| 3 | CC | M | 0.197 | 0.163 | 0.237 |
| 3 | S1BF | M | 0.227 | 0.192 | 0.265 |
| 7 | CC | M | 0.333 | 0.297 | 0.372 |
| 7 | S1BF | M | 0.376 | 0.338 | 0.416 |
